# Characterization of Cellular Senescence in Primary Human Astrocytes

**DOI:** 10.64898/2026.04.29.721581

**Authors:** Trenton A. Woodham, Maxfield M.G. Kelsey, John M. Sedivy

## Abstract

Senescent astrocytes have been identified in the brains of patients with neurodegenerative disorders, including Alzheimer’s disease, yet the molecular characteristics of replicative senescence in human astrocytes remain largely unexplored. Prior work has been hampered by the low proliferative capacity and limited telomere shortening of primary human astrocytes in culture. Here, we describe a culture system in which primary human astrocytes propagated under physiological (3%) oxygen reach canonical telomeric replicative senescence after extensive expansion (up to ∼76 population doublings). Senescence was confirmed through multiple biomarkers, including reduced EdU incorporation, elevated senescence-associated beta-galactosidase (SA-β-gal) activity, persistent DNA damage foci (γH2AX and 53BP1) predominantly localized to telomeres, and nuclear accumulation of p53.

RNA sequencing across a 12-week time course revealed early upregulation of young LINE-1 (L1HS) retrotransposon transcripts, type-I interferon (IFN-I) and senescence-associated secretory phenotype (SASP) pathway genes, alongside downregulation of cell-cycle and DNA repair programs. To resolve L1HS expression at individual locus resolution, we performed Nanopore DNA sequencing to generate a custom reference genome incorporating non-reference LINE-1 insertions. Applying our TE-Seq pipeline, we identified two full-length intergenic L1HS elements consistently upregulated across the replicative senescence time course, one of which, L1HS_9q22.32_2, retained intact ORF1 and ORF2 open reading frames, indicating potential retrotransposition competence.

To contextualize the astrocyte replicative senescence program, we compared it to three additional conditions. First, parallel astrocyte cultures maintained under normoxic (20%) oxygen entered senescence earlier and showed stronger SASP upregulation. Second, DNA damage-induced senescence (DDIS) triggered by etoposide treatment produced a stronger pro-inflammatory transcriptional signature than replicative senescence, including elevated IL6, IL1A, and IL1B expression. DDIS also upregulated L1HS_9q22.32_2 as well as a second intact element, L1HS_14q23.2_3, which we have previously identified among the small number of intact L1HS loci activated during replicative senescence in fibroblasts. The convergent activation of these intact elements across cell types and senescence modalities reinforces L1HS-driven IFN-I signaling as a conserved feature of the senescent program. Third, comparison with replicatively senescent fibroblasts revealed cell-type-specific SASP regulation: the pro-inflammatory cytokines IL6 and CCL2 were downregulated in senescent astrocytes relative to proliferating cells, opposite to their behavior in fibroblasts.

Together, these data establish the first comprehensive transcriptomic profile of replicative senescence in human astrocytes, offering a resource for understanding brain aging and senescence-associated neurodegeneration.

## Introduction

Aging is a complex biological process driven by multiple interacting hallmarks, one of which is cellular senescence, the permanent exit of a cell from the cell cycle [1]. Senescent cells remain metabolically active and can influence their tissue environment through the senescence-associated secretory phenotype (SASP), a program of secreted pro-inflammatory cytokines, growth factors, and matrix-remodeling enzymes [2]. The SASP contributes to chronic, low-grade inflammation and has been linked to tissue dysfunction in aged organisms [3].

Astrocytes are the most abundant cell type in the brain and serve critical roles in maintaining the blood-brain barrier, regulating neuronal metabolism and signal transduction, and responding to injury through reactive astrogliosis [4]. Although normally quiescent in the adult brain, astrocytes can become reactive and proliferative in response to injury or disease, forming glial scars essential for axonal regrowth through damaged regions [5, 6]. This proliferative capacity raises the question of whether astrocytes can undergo replicative senescence. Evidence from neurodegenerative disease strongly suggests they can. In Alzheimer’s disease, astrocytes in aged and AD-positive brains show elevated p16^INK4a^ alongside increased expression of the SASP factor MMP-1 and senescence-associated beta-galactosidase, with levels further increased in AD relative to age-matched healthy donors—a pattern consistent with astrocyte senescence contributing to both normal brain aging and disease pathology [7]. In a Parkinson’s disease model, the oxidative stressor paraquat induced astrocyte senescence both in vitro and in vivo, as measured by p16^INK4a^ expression and DNA damage markers, and depletion of these senescent astrocytes protected mice from developing associated neuropathology [8]. In amyotrophic lateral sclerosis, a rat model showed astrocytes acquiring a senescent phenotype at an accelerated rate compared to non-diseased controls, and this premature senescence was reversible [9].

While these studies have provided valuable insight into astrocyte senescence in disease contexts, the transcriptional landscape of replicative senescence in human astrocytes has not been characterized. Replicative senescence, driven by telomere attrition during successive cell divisions, is considered the most physiologically relevant model of senescence [10, 11], and because different senescence inducers can produce distinct downstream phenotypes, including differences in SASP composition [2, 12, 13], characterizing replicative senescence specifically is important. Here, we cultured primary human astrocytes to replicative exhaustion, confirmed senescence through canonical biomarkers, and performed RNA sequencing to define the global transcriptional changes associated with this process. We examined senescence development over a 12-week time course, the effects of culture oxygen conditions on the senescent transcriptome, and transcriptional differences between replicative and DNA-damage-induced senescence, and compared our astrocyte findings with published data from replicatively senescent human fibroblasts. Our results provide the first comprehensive transcriptomic characterization of replicative senescence in human astrocytes, revealing both conserved and cell-type-specific features of the senescent program.

## Results

### Astrocytes undergo replicative senescence in vitro

Primary human astrocytes cultured in both normoxic (20% O_2_) and physiological (3% O_2_) oxygen conditions exhibited a finite number of subcultures, with more subcultures generated under physiological oxygen (Figure 1A). Under physiological oxygen, astrocytes proliferative lifespan (37 passages, ∼76 population doublings), rivals that of classical fibroblast models. Proliferation slowed progressively in later passages, and late-passage astrocytes adopted an enlarged, flattened morphology characteristic of senescent cells (Figure 1B). EdU incorporation assays revealed that over 60% of early-passage astrocytes underwent S phase during a 48-hour labeling period, while fewer than 20% of final-passage cells incorporated EdU, with this fraction decreasing further over time (Figure 1C, D). Cells were designated as arrested and entering senescence 28 days after the final passage based on EdU incorporation levels.

**Figure 1:**
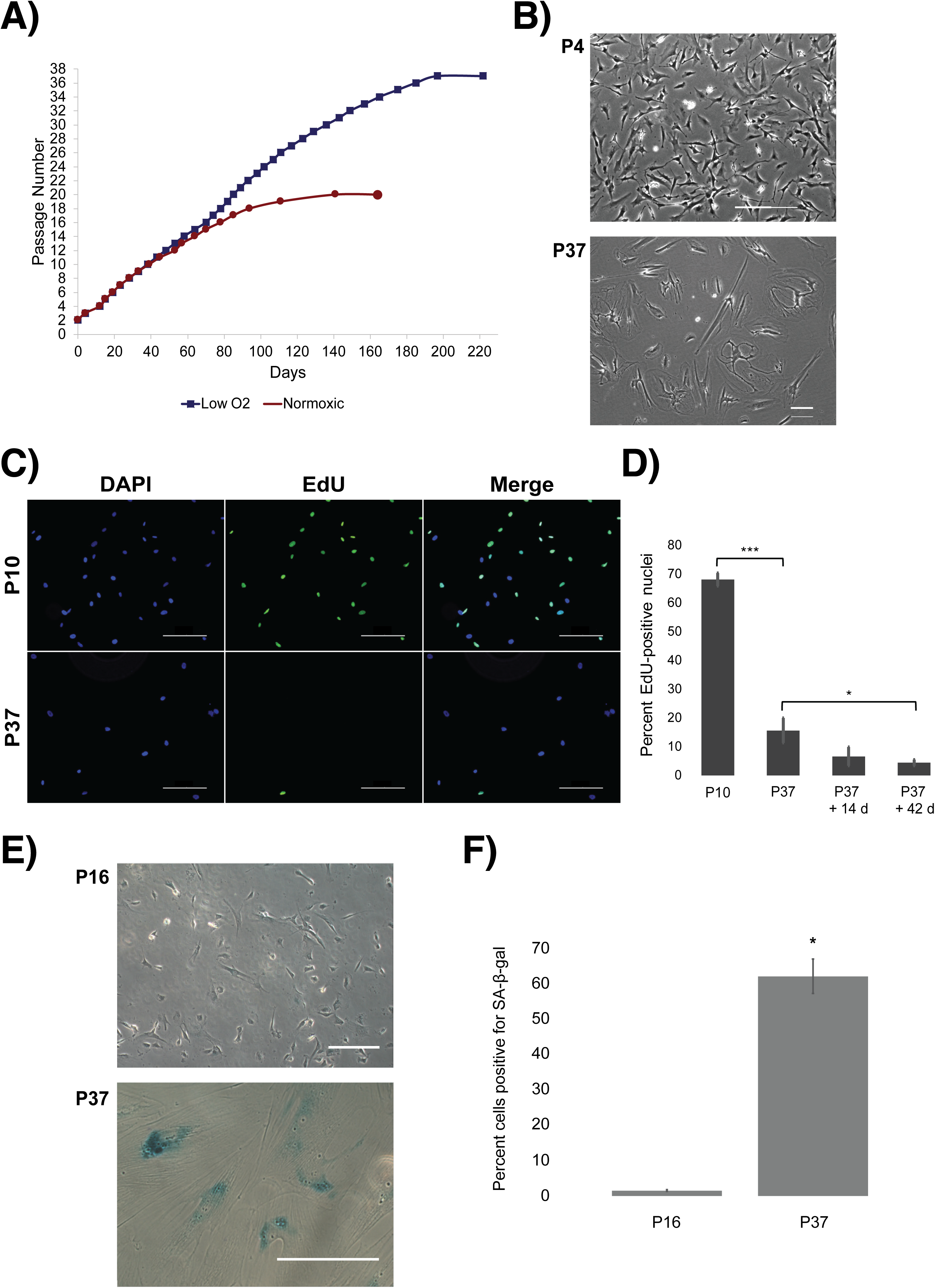
Primary human astrocytes have a limited growth span in culture. A) Astrocytes were grown in normoxic and physiological oxygen conditions. Passaging was performed on a culture at 90% confluence, and cultures split at a ratio of 1:4. Each passage is approximately 2 population doublings. B) Representative images of morphological changes in early (P4) and late (P37) passage astrocytes observed and recorded using phase contrast microscopy. Scale bars 100 µm. C) Representative images of early (P10) and late (P37) passage astrocytes treated with and stained for EdU. Imaged using fluorescent microscopy at 10x objective. Scale bars 200 µm. D) Fluorescent images were scored by counting total cells (DAPI) and positive cells (EdU) to calculate the percent positive number. Cells scored: 489 for P10, 189 for P37, 163 for P37 + 14 days, 378 for P37 + 42 days. P-values calculated via student t-test: *** < 1e-11, * < 0.02. E) Representative images of early passage (P16) and late passage (P37) human astrocytes after SA-β-gal staining. Fields were imaged using a color camera in both brightfield and phase contrast, then overlaid to show both the cell outline and stain. Scale bars 200 µm. F) Images were scored by counting total cells and cells with stain development, in order to calculate percent positivity. P16: 1,586 cells scored; P37: 140 cells scored. Statistics performed using a student t-test. P-value < 1e-5.

SA-β-gal activity was significantly elevated in late-passage (P37) astrocytes compared to early-passage (P16) controls. Approximately 62% of P37 astrocytes stained positive for SA-β-gal activity at pH 6.0, compared to 1.5% of P16 cells (Figure 1E, F). Immunofluorescence for the DNA damage markers γH2AX and 53BP1 showed significantly more damage foci in replicatively senescent (P37) and DNA-damage-induced senescent (DDIS) astrocytes than in proliferative controls (P12), with both higher percentages of positive nuclei and greater numbers of foci per nucleus in senescent conditions (Figure 2A–C, Supplemental Figure 2). Confocal imaging of p53 revealed diffuse cytoplasmic signal in proliferative astrocytes, nuclear localization in acutely damaged and newly arrested (P37) cells, and diminished, predominantly cytoplasmic signal in cells that had been senescent for 5 weeks, consistent with p53 activation during senescence entry followed by subsequent inactivation (Supplemental Figure 3). Together, these results demonstrate that cultured human astrocytes undergo replicative senescence with hallmarks consistent with senescence in other cell types.

**Figure 2:**
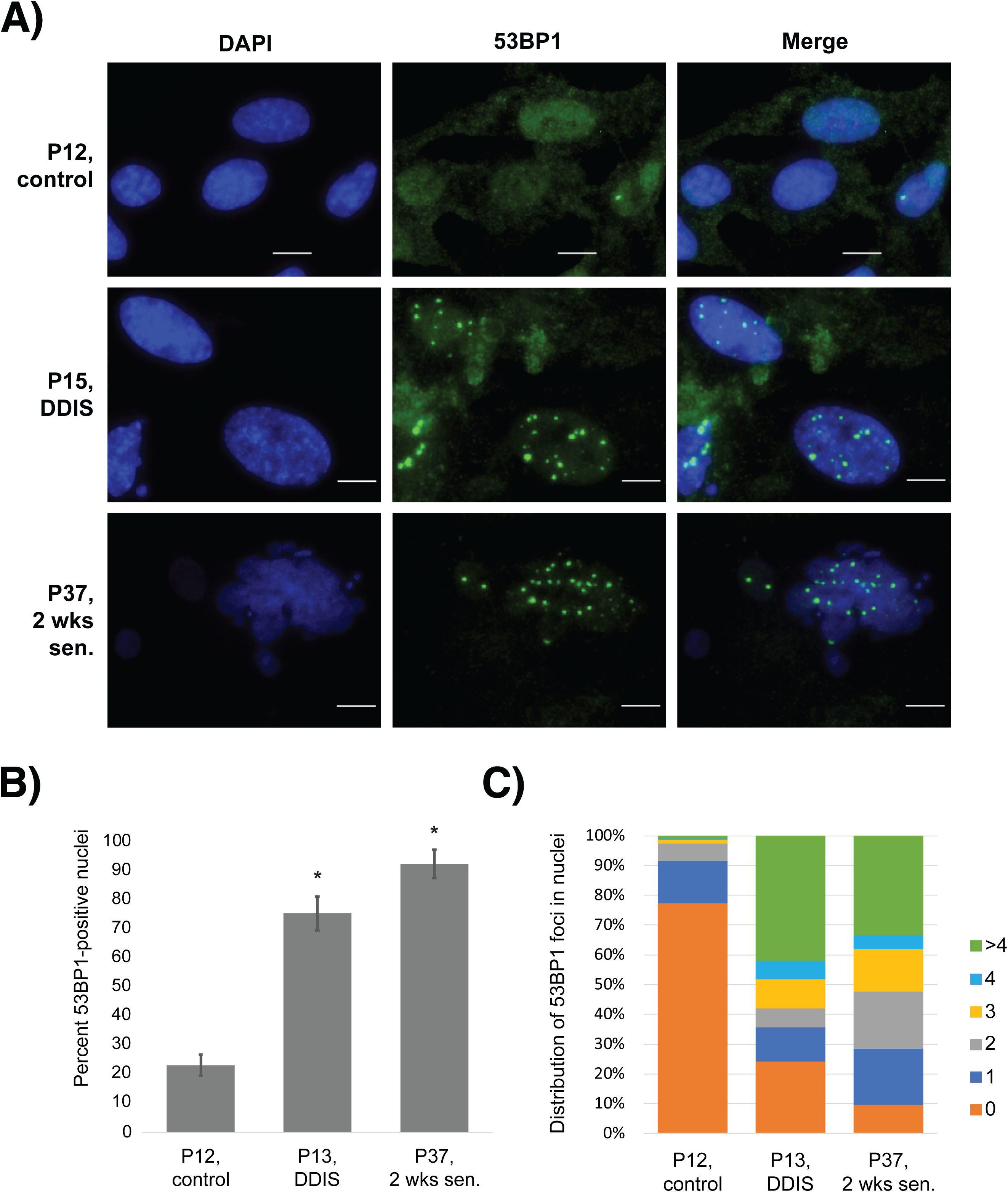
53BP1 immunofluorescent staining. A) Conditions imaged were early proliferative (P12), DNA-damaged (P15, DDIS), and early senescent (P37, 2 wks sen) astrocytes. Scale bars 10 µm. B) Percent positivity calculated by tabulating total nuclei and number of nuclei with at least 1 damage focus co-staining in nuclear region. Statistics performed using student t-test. P-value: * < 0.0001. C) Distribution of damage foci in cells. Cells were scored by counting the number of foci per nucleus and sorting the cells into groups by number of foci. P12, control: 336 cells scored; P13, DDIS: 112 cells scored; P37, 2 wks sen: 22 cells scored.

### The transcriptomic landscape of replicative senescence in human astrocytes

To characterize the transcriptomic landscape of astrocyte senescence, we performed RNA-seq on astrocytes at five stages: early passage (EP), senescence entry (Sen0), and 4, 8, and 12 weeks post-arrest (Sen4, Sen8, Sen12), all cultured under physiological oxygen. We also sequenced astrocytes cultured under normoxic (20% O_2_) conditions and astrocytes undergoing DNA damage-induced senescence (DDIS). Principal component analysis separated proliferating from senescent samples along PC1 (69% of variance), and we observed tight clustering among replicates (Supplemental Figure 1A,B).

Differential expression analysis revealed extensive transcriptional remodeling upon senescence entry, with 9,377 significantly regulated genes (≥2-fold, FDR < 0.05) in Sen0 vs. EP and 8,434 in Sen12 vs. EP. The majority of these changes were shared: 3,865 genes were upregulated and 3,401 downregulated at both time points (Supplemental Figure 4A, B). Direct comparison of Sen12 to Sen0 yielded only 843 significantly regulated genes, and GSEA revealed no KEGG pathways significantly different between the two stages, indicating that the overall pathway architecture of senescence is largely established at entry and remains stable thereafter.

GSEA using KEGG pathways confirmed upregulation of immune response programs (IFN-I, SASP, complement and coagulation cascades) and cytochrome P450/drug metabolism pathways, alongside downregulation of cell cycle, DNA replication, and DNA repair pathways (homologous recombination, mismatch repair, nucleotide excision repair) (Figure 3A–C). The IFN-I and SASP pathways were significantly enriched at both Sen0 and Sen12, though the SASP pathway did not reach statistical significance by GSEA at Sen0. However, hierarchical clustering of SASP gene expression (Figure 4) confirmed that SASP upregulation is evident at Sen0: EP samples clustered separately from all senescent time points, confirming a broad shift in secretory gene expression upon senescence entry. A large subset of SASP genes, including chemokines (CCL1, CCL7, CCL20, CXCL3), growth factors (FGF2, EREG, FGF7, WNT2), and signaling components (AXL, CD9, AREG), was consistently upregulated across all senescent time points relative to EP. Other SASP components showed a more dynamic pattern, with expression increasing progressively through the time course, reaching their highest levels only at Sen8 or Sen12 (e.g., CXCL1, CXCL8, IGFBP1, IGFBP2). However, several canonical pro-inflammatory SASP factors behaved opposite to their well-characterized pattern in fibroblasts: IL6 and CCL2 had higher relative expression in EP than in senescent conditions, and IL1A was similarly downregulated in Sen12 relative to EP [3, 14, 15]. This may reflect a role for these cytokines in reactive, proliferating astrocytes, where IL6 is a known modulator of reactive astrogliosis through STAT3 signaling [16]. Because existing SASP gene sets are derived largely from work in fibroblasts, these results suggest that while astrocytes engage a substantial portion of the canonical secretory program upon senescence, they also diverge in important ways, mounting a qualitatively distinct SASP rather than simply a weaker version of the fibroblast response. IFN-I pathway genes were broadly upregulated in senescent conditions, though IFNA1, typically upregulated in senescent fibroblasts [3], was more highly expressed in EP astrocytes than in most senescent samples (Figure 4).

**Figure 3:**
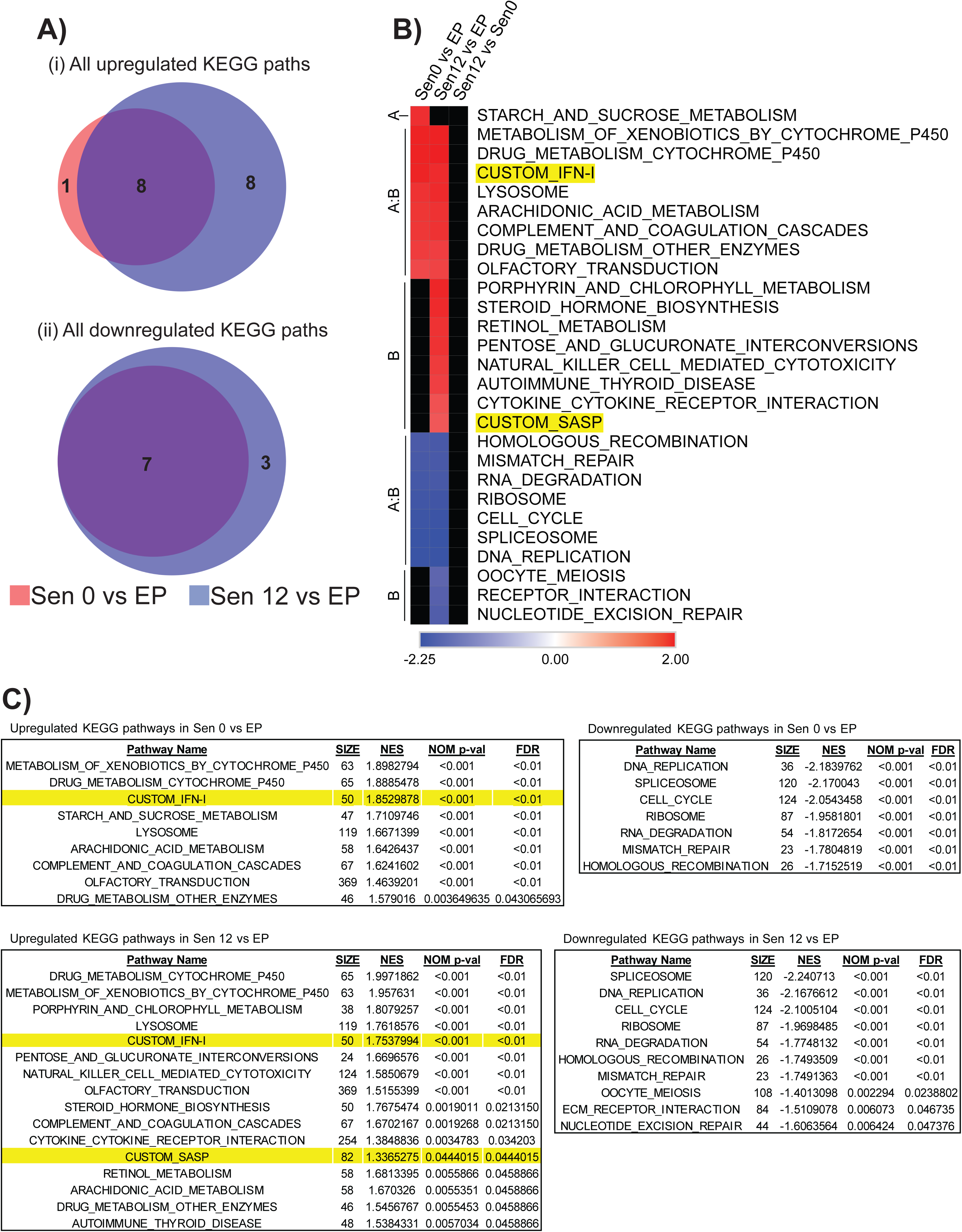
Pathway-level differences in time points of replicative senescence. A) Overlap of upregulated (ii) and downregulated (ii) KEGG pathways in Sen0 vs. EP and Sen12 vs. EP conditions. FDR cutoff 0.05. B) Heatmap of significantly regulated KEGG pathways in comparisons Sen0 vs. EP (Group A) and Sen12 vs. EP (Group B). Legend represents normalized enrichment scores (NES). FDR cutoff 0.05. Black regions indicate pathways that did not meet the FDR cutoff. Highlighted pathways were manually added to the analysis and are not standard KEGG pathways. C) Tables for regulated KEGG pathways in Sen0 vs. EP and Sen12 vs. EP conditions. Highlighted pathways were manually added to the analysis and are not standard KEGG pathways. FDR cutoff 0.05. “SIZE”: total number of genes in pathway; “NES”: normalized enrichment score; “NOM p-val”: p-value calculated by GSEA; “FDR”: corrected false discovery rate by performing Benjamini-Hochberg procedure on NOM p-values.

**Figure 4:**
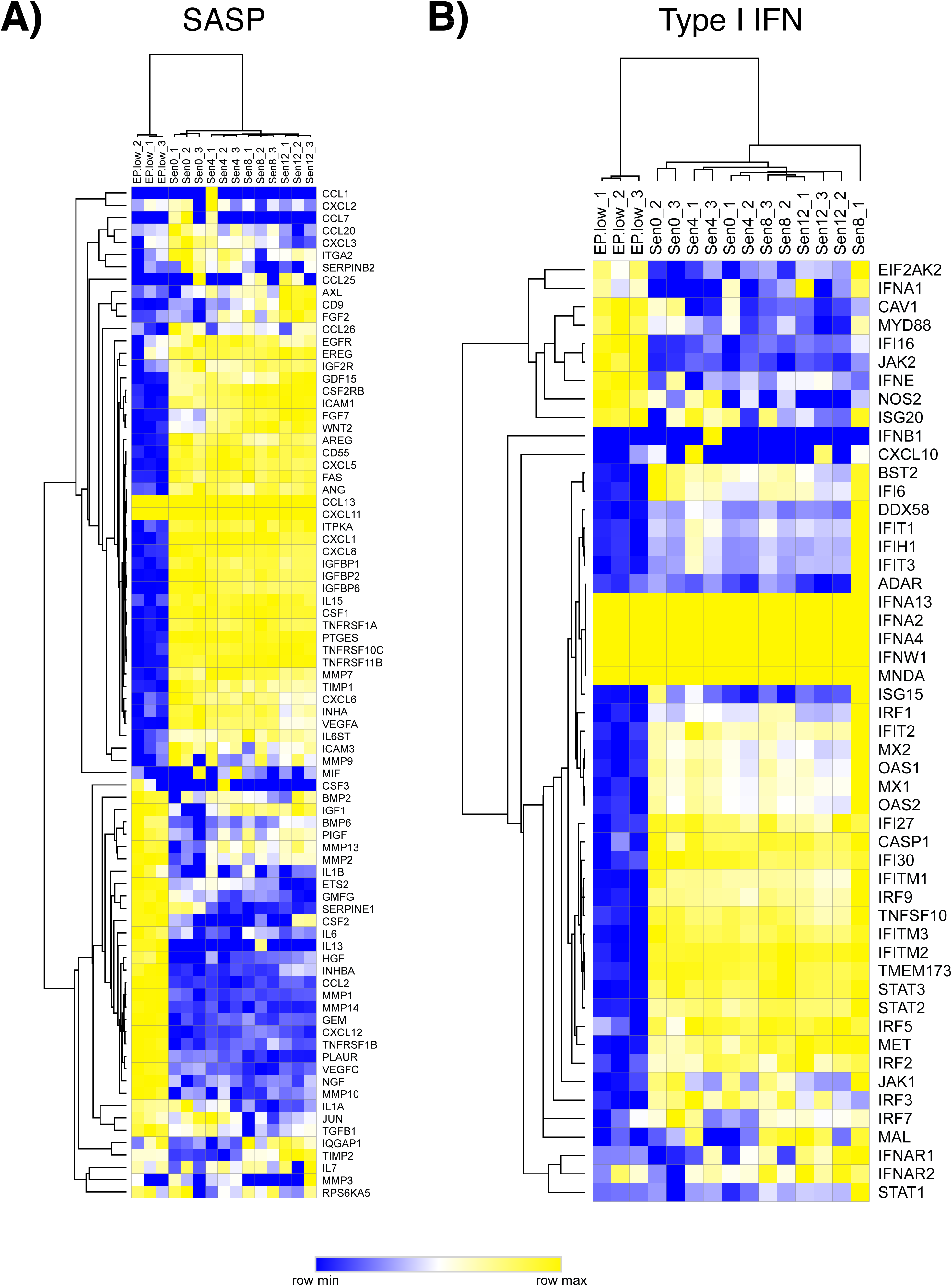
SASP and IFN-I gene expression during astrocyte replicative senescence. Heatmap displaying relative expression behavior of genes in the SASP (A) and IFN-I (B) gene list for each sample and time point. Hierarchical clustering performed for both rows and columns using a one minus pearson correlation metric. For genes where every sample has the same expression value, entire row is represented as “row max”.

### Oxygen culture conditions affect the senescent astrocyte transcriptome

Standard culture conditions use atmospheric oxygen levels (20%), but oxygen levels within non-lung tissues typically range from 2–4% [17]. To assess how oxygen affects astrocyte senescence, we cultured the same lot of astrocytes in parallel under physiological (3% O_2_) and normoxic (20% O_2_) conditions. Astrocytes in normoxic conditions senesced earlier than those cultured at physiological oxygen (Figure 1A). At the pathway level, the IFN-I pathway was upregulated only under physiological oxygen, while the SASP pathway was upregulated only under normoxic conditions (Figure 5B). Direct comparison of Sen0-atm vs. Sen0-low showed upregulation of the SASP, cytokine interaction, and protein export/ubiquitination pathways in normoxic senescence (Figure 5C). Within the SASP pathway, IL6 and CCL2 were downregulated in physiological-oxygen senescence but upregulated in normoxic senescence relative to EP (Supplemental Figure 5D). These results demonstrate that oxygen conditions substantially influence the transcriptional profile of senescent astrocytes and that standard normoxic culture produces a distinct senescence program.

**Figure 5:**
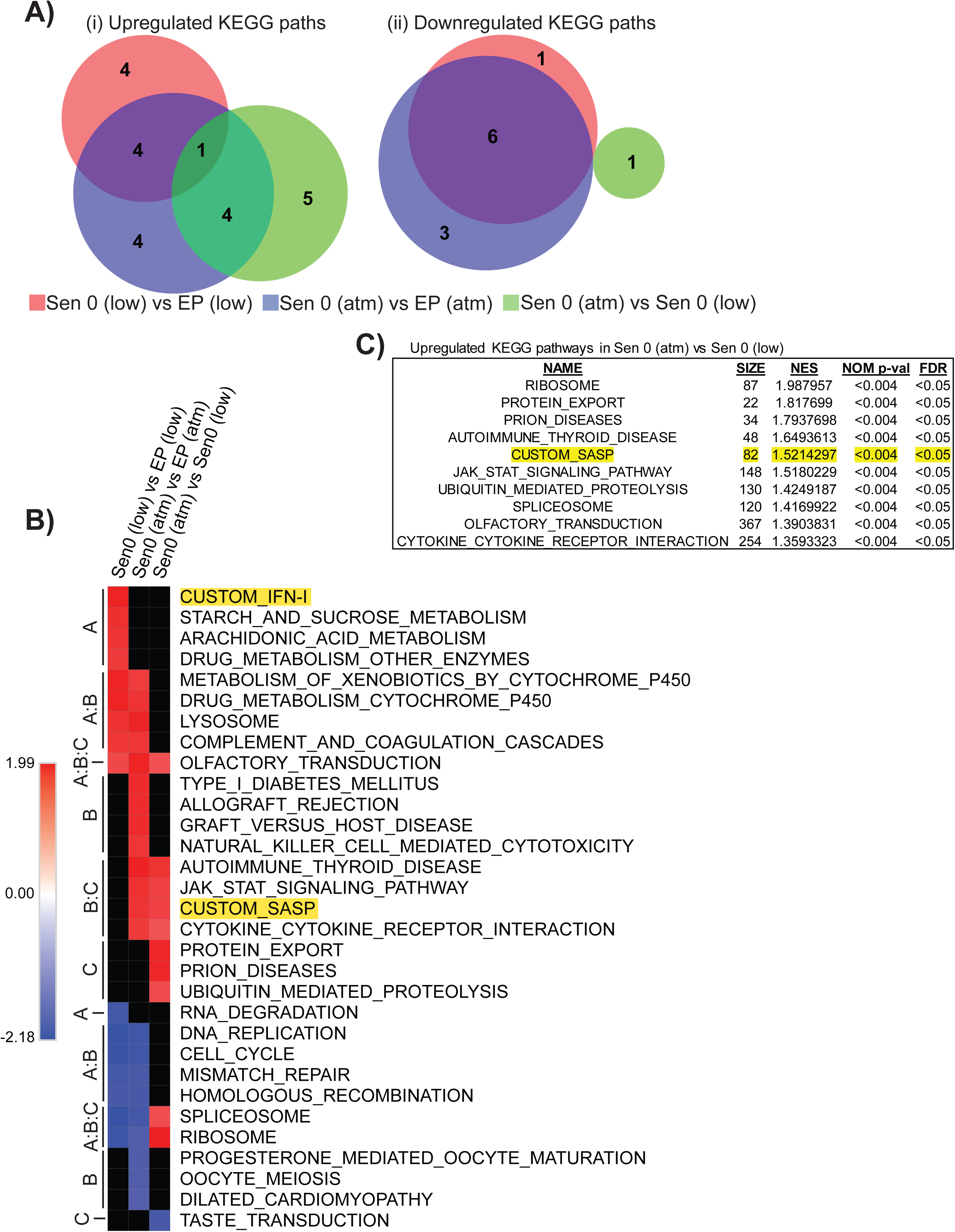
Pathway-level differences in replicatively senescent astrocytes from varying oxygen culture conditions. D) Overlap of upregulated (ii) and downregulated (ii) KEGG pathways in Sen 0 (low) vs. EP (low) [red], Sen 0 (atm) vs. EP (atm) [blue], and Sen0 (atm) vs. Sen 0 (low) [green] conditions. FDR cutoff 0.05. E) Heatmap of significantly regulated KEGG pathways in comparisons Sen 0 (low) vs. EP (low) [Group A], Sen 0 (atm) vs. EP (atm) [Group B], and Sen 0 (atm) vs. Sen 0 (low) [Group C]. Legend represents normalized enrichment scores (NES). FDR cutoff 0.05. Black regions indicate pathways that did not meet the FDR cutoff. Highlighted pathways were manually added to the analysis and are not standard KEGG pathways. F) Table of upregulated KEGG pathways in Sen 0 (atm) vs. Sen 0 (low). Highlighted pathways were manually added to the analysis and are not standard KEGG pathways. FDR cutoff 0.05. “SIZE”: total number of genes in pathway; “NES”: normalized enrichment score; “NOM p-val”: p-value calculated by GSEA; “FDR”: corrected false discovery rate by performing Benjamini-Hochberg procedure on NOM p-values.

### DNA-damage-induced senescence produces a stronger pro-inflammatory signature than replicative senescence

To compare replicative senescence with another common senescence model, we induced DNA-damage-induced senescence (DDIS) in astrocytes using etoposide (20 µM, 72 hours) and allowed senescence to develop over 8 weeks before RNA-seq. DDIS showed more significantly regulated KEGG pathways than 12-week replicative senescence (24 upregulated and 17 downregulated vs. 16 and 10, respectively; Figure 6A). The two models shared upregulation of drug metabolism, immune response, SASP, and IFN-I pathways and downregulation of cell cycle and DNA repair, but DDIS uniquely showed significant regulation of JAK-STAT signaling and additional transcription-related pathways (Figure 6B). Direct comparison of DDIS vs. Sen12 revealed upregulation of ECM receptor interaction and cytokine-cytokine interaction pathways in DDIS.

**Figure 6:**
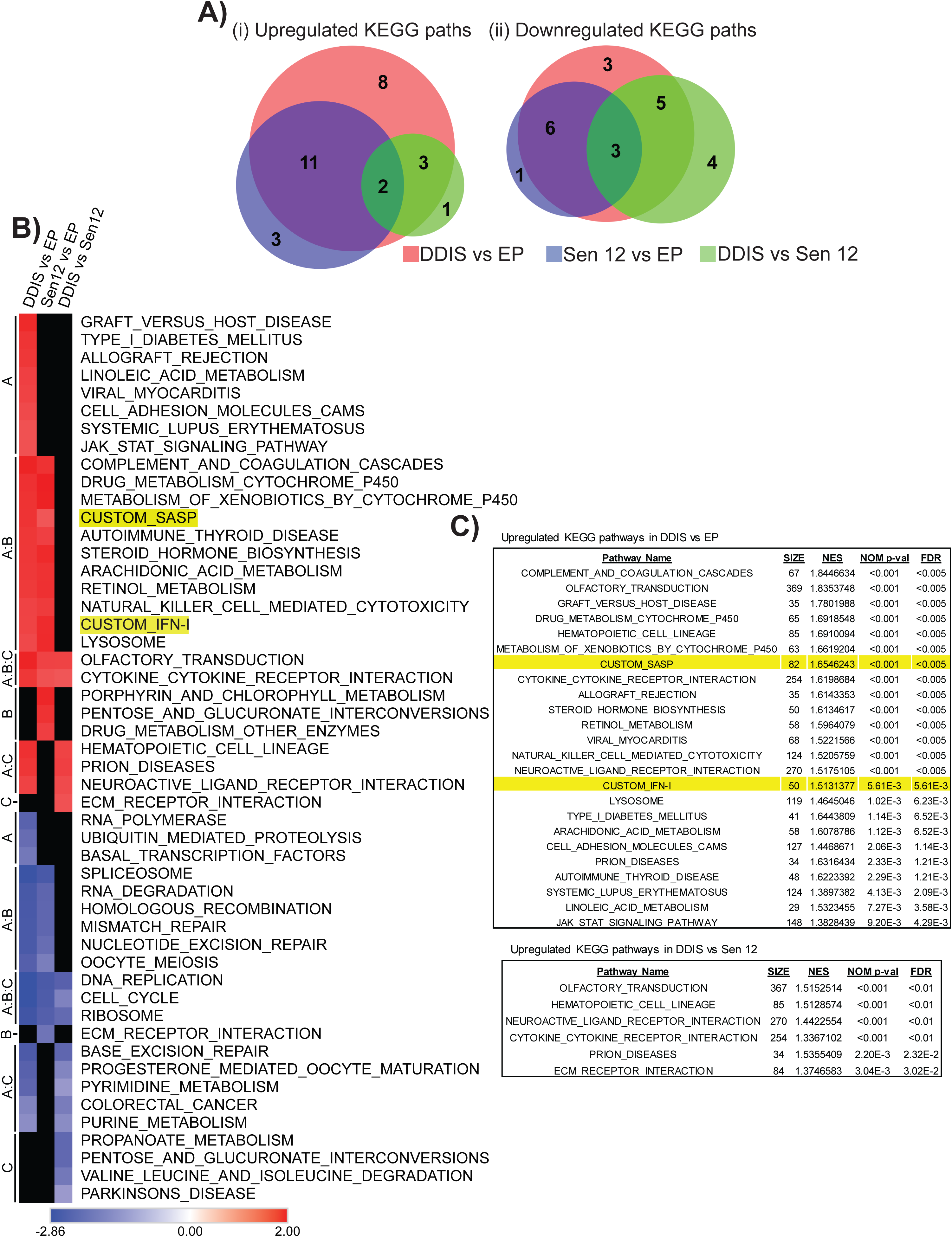
Pathway level differences in DDIS compared to replicative senescence in astrocytes. G) Overlap of upregulated (i) and downregulated (ii) KEGG pathways in DDIS vs. EP [red], Sen12 vs. EP [blue], and DDIS vs. Sen12 [green] conditions. FDR cutoff 0.05. H) Heatmap of significantly regulated KEGG pathways in comparisons DDIS vs. EP [Group A], Sen12 vs. EP [Group B], and DDIS vs. Sen12 [Group C]. Legend represents normalized enrichment scores (NES). FDR cutoff 0.05. Black regions indicate pathways that did not meet the FDR cutoff. Highlighted pathways were manually added to the analysis and are not standard KEGG pathways. I) Tables for upregulated KEGG pathways in DDIS vs. EP and DDIS vs. Sen12 conditions. Highlighted pathways were manually added to the analysis and are not standard KEGG pathways. FDR cutoff 0.05. “SIZE”: total number of genes in pathway; “NES”: normalized enrichment score; “NOM p-val”: p-value calculated by GSEA; “FDR”: corrected false discovery rate by performing Benjamini-Hochberg procedure on NOM p-values.

Notable differences were apparent at the gene level within the SASP pathway: IL1A, IL1B, and IL6 were upregulated in DDIS vs. EP but downregulated in Sen12 vs. EP, and were significantly higher in DDIS than Sen12 when compared directly (Supplemental Figure 6D). Chemokines and growth factors (VEGFA, FGF2) were also uniquely upregulated in DDIS. Within the IFN-I pathway, IRF7 was upregulated in DDIS but not in Sen12 relative to EP, while IFNE was downregulated in Sen12 but upregulated in DDIS (Supplemental Figure 6C). Together, these data indicate that DDIS produces a substantially stronger pro-inflammatory transcriptional signature than replicative senescence in astrocytes.

### Astrocyte replicative senescence shares core pathways with fibroblasts but features a less pro-inflammatory profile

To contextualize our findings, we compared transcriptomic data from 12-week replicatively senescent astrocytes (Sen12) with previously published data from 16-week replicatively senescent human fibroblasts (Sen16) [3]. At the pathway level, most KEGG pathways regulated in astrocyte senescence were also regulated in fibroblast senescence, including shared upregulation of IFN-I, SASP, complement cascades, and cytochrome P450 pathways, and shared downregulation of cell cycle and DNA repair (Figure 7A, B).

**Figure 7:**
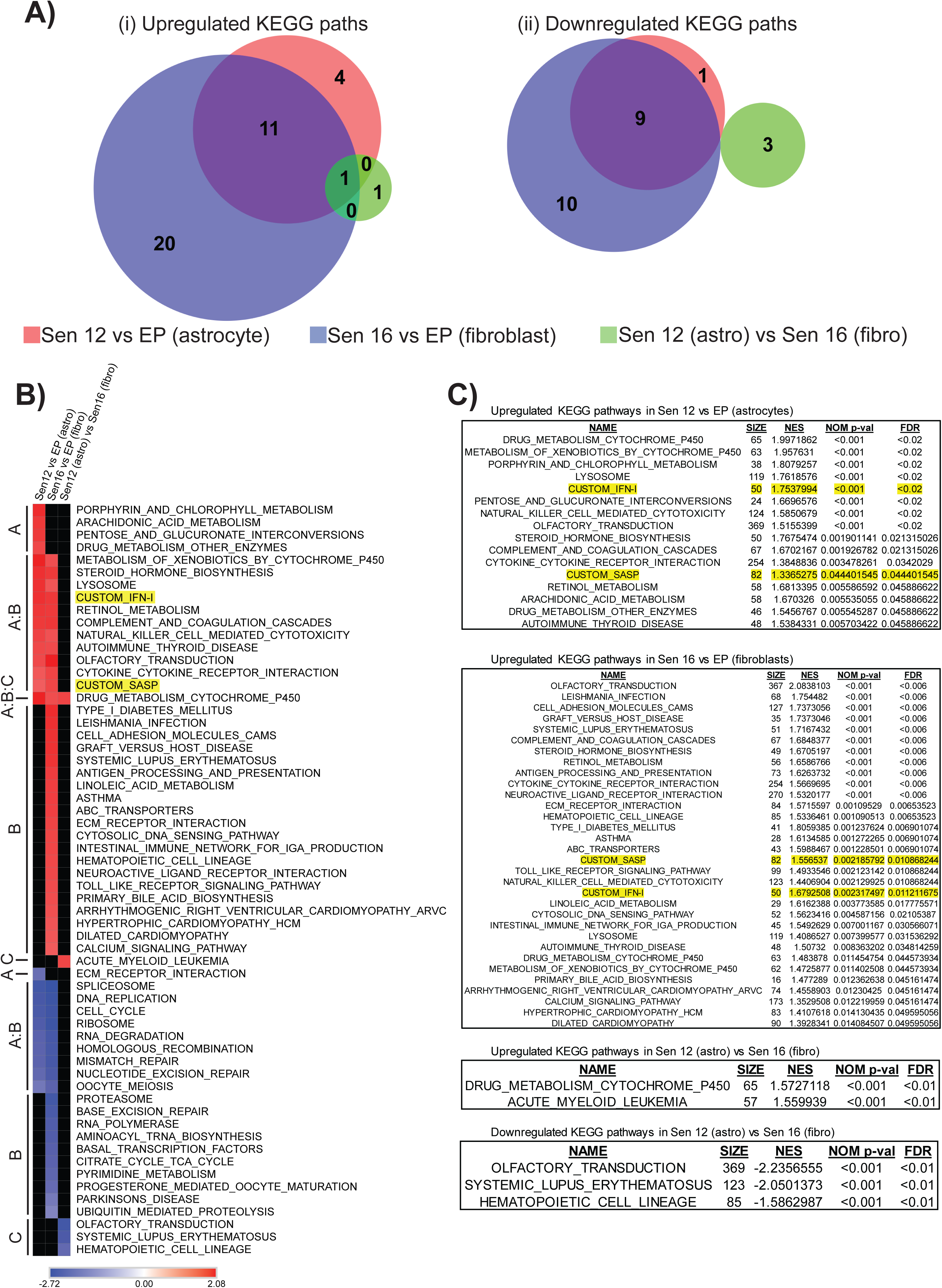
Pathway level expression changes in replicative senescence modeled in astrocytes and fibroblasts. A) Overlap of upregulated (i) and downregulated (ii) KEGG pathways in Sen12 vs. EP (astrocytes) [red], Sen16 vs. EP (fibroblasts) [blue], and Sen12 (astro) vs. Sen16 (fibro) [green] conditions. FDR cutoff 0.05. B) Heatmap of significantly regulated KEGG pathways in Sen12 vs. EP (astro) [Group A], Sen16 vs. EP (fibro) [Group B], and Sen12 (astro) vs. Sen16 (fibro) [Group C]. Legend represents normalized enrichment scores (NES). FDR cutoff 0.05. Black regions indicate pathways that did not meet the FDR cutoff. Highlighted pathways were manually added to the analysis and are not standard KEGG pathways. C) Tables for upregulated KEGG pathways in Sen12 vs. EP (astrocytes), upregulated pathways in Sen16 vs. EP (fibroblasts), and up- and down-regulated pathways in Sen12 (astro) vs. Sen16 (fibro). Highlighted pathways were manually added to the analysis and are not standard KEGG pathways. FDR cutoff 0.05. “SIZE”: total number of genes in pathway; “NES”: normalized enrichment score; “NOM p-value”: p-value calculated by GSEA; “FDR”: corrected false discovery rate by performing Benjamini-Hochberg procedure on NOM p-values.

Cell-type differences were apparent in both the IFN-I and SASP pathways. Within the IFN-I pathway, interferon-alpha genes (IFNA1, IFNA13) and IFNE were upregulated in senescent fibroblasts but unchanged or downregulated in senescent astrocytes (Figure 7C). The SASP pathway presented a more complex picture. Several canonical pro-inflammatory SASP factors, including IL1A, IL1B, IL6, CCL2, MMP3, and IGF1, were upregulated in senescent fibroblasts but downregulated or unchanged in senescent astrocytes (Figure 7D). However, other SASP components were regulated in the opposite direction, with genes such as WNT2, EREG, IL15, and CCL26 upregulated in senescent astrocytes but not in fibroblasts. These findings indicate that while the core senescence program is conserved between cell types, the composition of the SASP diverges substantially: astrocytes do not simply mount a weaker secretory response, but rather a qualitatively distinct one, with potential implications for how senescent astrocytes influence the brain microenvironment compared to senescent fibroblasts in peripheral tissues.

### LINE-1 retrotransposon expression in senescent astrocytes

RT-qPCR analysis showed upregulation of LINE-1 (L1HS) retrotransposon transcripts in both replicatively senescent and DDIS astrocytes, with the 5’ UTR amplicon elevated approximately 2-fold in replicative senescence and 2.5-fold in DDIS relative to proliferative cells (Figure 8A), consistent with LINE-1 activation during senescence-associated chromatin remodeling as previously reported in fibroblasts [3, 15, 16].

**Figure 8:**
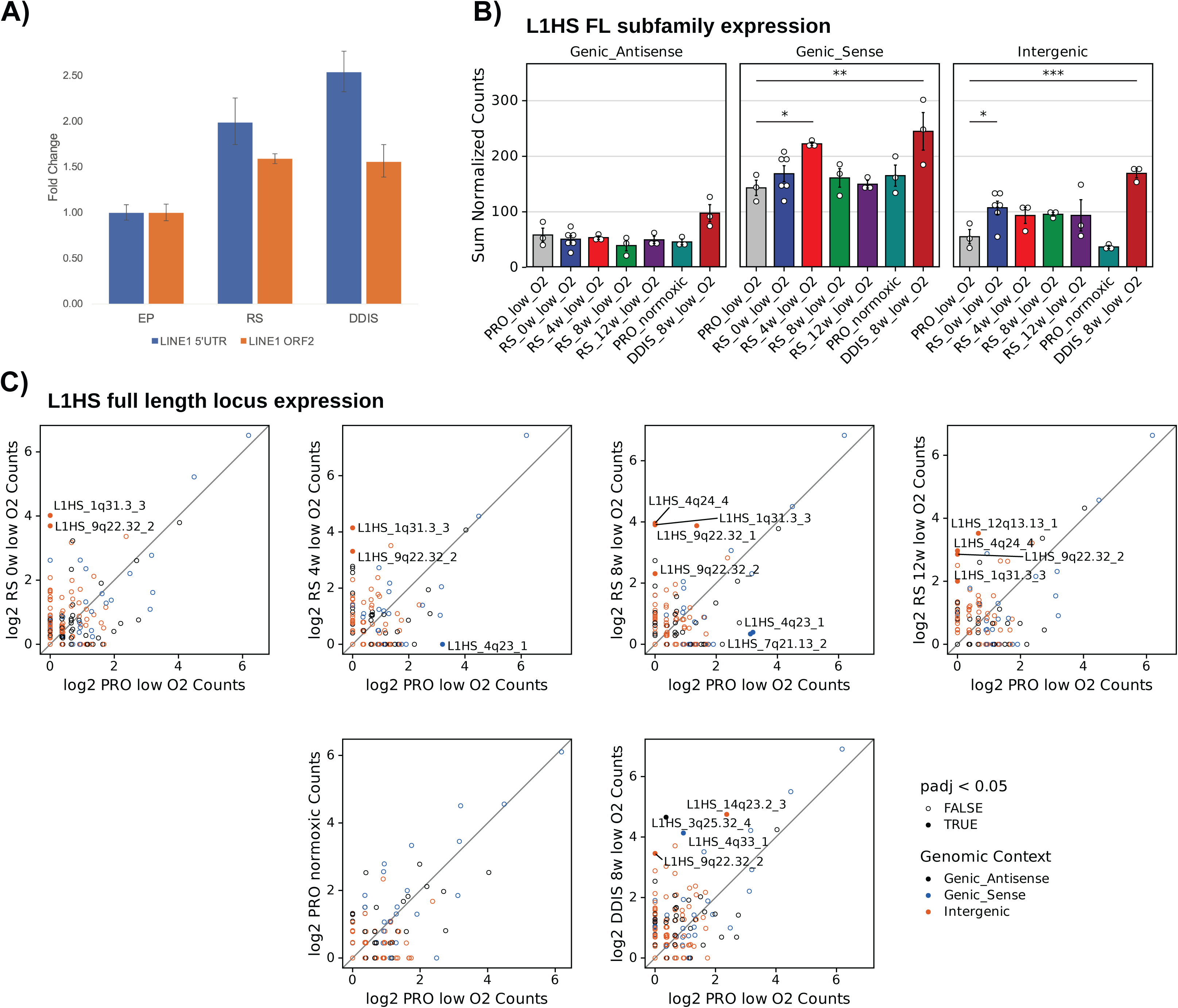
L*I*NE*-1* transcripts are more highly expressed in senescent than proliferative astrocytes. A) EP—early passage; RS—replicative senescence; DDIS—DNA-damage induced senescence. Blue bars—5’ UTR primers; orange bars—ORF2 primers. 1 sample per condition tested in quadruplet. Normalized to GAPDH. Fold-change calculated through 2-ΔΔCT method. Error bars represent absolute values of 2-ΔΔCT relative to sample added to the associated +/- standard deviations. B) Bar plots showing total DESeq2-normalized read counts from various categories of L1HS elements. Statistical comparisons are based on negative binomial modeling of raw aggregated counts, incorporating DESeq2-derived size factors as a model offset; p-values are FDR-adjusted for all pairwise comparisons in the panel. FDR-adjusted p ≤ 0.05 *, ≤ 0.01 **, and ≤ 0.001 ***. C) Scatter plot showing log2 DESeq2 normalized counts (+ 1) of every full-length L1HS element, colored by genomic context. Differentially expressed elements (FDR ≤ 0.05) are denoted by filled in circles and are labeled by their ID.

We next sought to characterize L1HS expression at the individual locus level using RNA-seq. Because many L1HS elements are polymorphic (present in some individuals but absent from the reference genome) standard alignment approaches fail to capture their expression. We therefore performed Nanopore long-read DNA sequencing of our astrocyte donor cells, generating 76.9 gigabases (∼25x genome coverage), and applied the TE-Seq pipeline [18] to construct a custom reference genome incorporating non-reference L1HS insertions and masking reference elements absent from the donor (Supplemental Figure 8). This TE-complete reference enabled accurate assignment of RNA-seq reads to individual L1HS loci, including polymorphic elements that would otherwise be invisible to expression analysis.

At the subfamily level, aggregated counts from full-length intergenic L1HS elements were significantly elevated in replicative senescence relative to proliferating cells (Figure 8B). Genic sense elements also showed significant upregulation, though since many L1HS elements reside within gene introns, this signal likely reflects passive co-transcription of host gene pre-mRNA rather than autonomous L1HS promoter activity. Genic antisense elements showed no significant change.

At the individual locus level, two intergenic elements, L1HS_1q31.3_3 and L1HS_9q22.32_2, were consistently upregulated across the replicative senescence time course (Figure 8C). L1HS_9q22.32_2 is an intact element, retaining full-length ORF1 and ORF2 open reading frames, and was also significantly upregulated in DDIS, suggesting convergent activation of this retrotransposition-competent element by distinct senescence triggers. DDIS additionally upregulated L1HS_14q23.2_3, another intact intergenic element (Figure 8C). Notably, we have previously identified L1HS_14q23.2_3 as one of only a small number of intact L1HS elements upregulated during replicative senescence in LF1 fibroblasts [19], indicating that this element is activated across both cell types and senescence modalities. The recurrent identification of intact, autonomously expressed L1HS elements in senescent astrocytes and fibroblasts is consistent with the view that retrotransposition-mediated DNA damage and cytoplasmic nucleic acid sensing contribute to the inflammatory phenotype of senescent cells, and suggests that L1HS activation in senescent astrocytes may be a source of sterile neuroinflammation relevant to age-associated neurodegeneration.

## Discussion

We present the first comprehensive characterization of replicative senescence in primary human astrocytes, demonstrating that these cells undergo a bona fide senescent program with conserved hallmarks—growth arrest, SA-β-gal activity, persistent DNA damage, and p53 activation (Figures 1–2, Supplemental Figure 3)— accompanied by widespread transcriptional changes, including a SASP and IFN-I response that is qualitatively distinct from that of fibroblasts.

Our RNA-seq time-course analysis revealed that the transcriptome is extensively remodeled upon senescence entry but remains largely stable thereafter: no KEGG pathways were significantly different between Sen0 and Sen12 (Figure 3), although 843 genes were significantly regulated between these time points. The IFN-I and SASP pathways were upregulated at senescence entry (Figure 3, 4).

Comparisons across senescence models and cell types revealed both conserved and context-dependent features of the senescent program. Astrocytes cultured under normoxic (20% O₂) conditions arrested earlier (Figure 1A) and displayed stronger SASP induction by GSEA than did cells grown under physiological (3% O₂) (Figure 5B, C). DNA damage-induced senescence produced a substantially stronger pro-inflammatory transcriptional signature than replicative senescence, with upregulation of IL1A, IL1B, IL6, and additional chemokines and growth factors (Figure 6, Supplemental Figure 6C, D), consistent with the view that exogenous DNA damage triggers a more acute SASP than telomere attrition-driven senescence [9]. Comparison with replicatively senescent fibroblasts revealed that while core senescence pathways are conserved, astrocytes mount a distinctly less pro-inflammatory response: the cytokines IL6 and CCL2 are downregulated rather than upregulated in replicatively senescent astrocytes relative to proliferative cells (Figure 7C, D). This reversal likely reflects the biology of reactive astrocytes: in culture, proliferating astrocytes are by definition in a reactive state, and IL6 is a known modulator of reactive astrogliosis through STAT3 signaling [16]. The association of IL6 and CCL2 with proliferating, reactive astrocytes rather than with senescence suggests that the astrocyte SASP has a fundamentally different composition from that of fibroblasts, with important implications for understanding how senescent astrocytes affect the brain microenvironment.

A key finding of this study is the identification of specific intact L1HS retrotransposon loci that are autonomously upregulated in senescent astrocytes. LINE-1 activation has been established as a feature of replicative senescence in fibroblasts, where de-repression of heterochromatic L1HS elements leads to accumulation of cytoplasmic reverse-transcribed DNA, activation of the cGAS-STING pathway, and induction of IFN-I signaling [3, 14, 15]. Our data demonstrate that this phenomenon extends to astrocytes: L1HS transcripts were elevated in both replicative and DNA damage-induced senescence by RT-qPCR (Figure 8A), and RNA-seq analysis of full-length intergenic L1HS elements confirmed upregulation at the subfamily level during replicative senescence (Figure 8B). Critically, by applying the TE-Seq pipeline with a Nanopore-derived custom reference genome, we were able to resolve this bulk signal to individual loci, identifying two intergenic L1HS elements (L1HS_1q31.3_3 and L1HS_9q22.32_2) consistently upregulated across the replicative senescence time course (Figure 8C). L1HS_9q22.32_2 retains intact ORF1 and ORF2 open reading frames, meaning it is potentially competent for retrotransposition. This element was also upregulated in DDIS, suggesting that its de-repression is a convergent feature of the senescent state regardless of the inducing stimulus. DDIS additionally activated L1HS_14q23.2_3, another intact intergenic element that we have previously identified as one of a small number of intact L1HS loci upregulated during replicative senescence in LF1 fibroblasts [19]. The activation of the same intact element across two cell types and multiple senescence modalities suggests that specific genomic loci may be recurrently vulnerable to senescence-associated chromatin remodeling, rather than L1HS de-repression occurring stochastically across the genome. The identification of retrotransposition-competent L1HS elements in senescent astrocytes has particular significance for the brain: astrocytes are the most abundant cell type in the CNS and are positioned to influence neuronal health through both paracrine signaling and direct metabolic support. If intact L1HS elements undergo retrotransposition in senescent astrocytes in vivo, the resulting DNA damage and innate immune activation could contribute to the sterile neuroinflammation that characterizes neurodegenerative diseases including Alzheimer’s disease. Consistent with this, IFN-I signaling, the downstream effector of cGAS-STING activation by cytoplasmic L1 DNA, was among the pathways most robustly upregulated across the senescence time course. Future work will be needed to determine whether L1HS retrotransposition occurs in senescent astrocytes and whether pharmacological inhibition of reverse transcriptase can attenuate the inflammatory phenotype, as has been demonstrated in senescent fibroblasts [3].

In summary, our work establishes a replicative senescence model for human astrocytes and provides a transcriptomic resource that reveals both conserved and astrocyte-specific features of the senescent program. The distinct SASP composition underscores the need to characterize senescence in cell types beyond fibroblasts, particularly those relevant to the aging brain, while the identification of specific intact L1HS loci activated across cell types and senescence modalities reinforces L1HS-driven IFN-I activation as a conserved feature of the senescent program.

## Materials and Methods

### Cell culture

Primary human astrocytes (ScienCell, Cat. 1800) were isolated from the cerebral cortex of a non-disease fetal donor, cryopreserved at passage 1, and verified GFAP-positive by the vendor. Cells were cultured at 37°C in either physiological oxygen (3% O_2_, 5% CO_2_) or normoxic (20% O_2_, 5% CO_2_) conditions in complete astrocyte medium (ScienCell, Cat. 1801: 2% FBS, 1% P/S, 1% AGS, 96% basal medium). Dishes (10-cm) were pre-treated with poly-L-lysine (PLL; 15 µg/ml). Astrocytes were seeded at 250,000 cells/dish, fed every 3 days with 100% medium exchange, and passaged at 1:4 ratio upon reaching 90% confluence (2 population doublings per passage). Passaging was ceased when cultures failed to reach 90% confluence within 3 weeks. Human lung fibroblasts (LF1) were cultured in physiological oxygen conditions in Ham’s F-10 medium supplemented with 15% FBS, 1% P/S, and 1% L-glutamine, and passaged at 1:4 ratio at 80% confluence.

### Population doubling calculations

At each passage, cells were counted using a hemocytometer. Population doubling levels (PDL) were calculated as: PDL = 3.32 × [log(n1) − log(n2)] + X, where n1 = cells harvested, n2 = cells seeded, and X = previous PDL.

### EdU incorporation assay

Astrocytes on PLL-coated coverslips were incubated with 10 µM EdU (Click-iT EdU Alexa Fluor 488 Imaging Kit, Invitrogen C10337) for 48 hours, fixed with 3.7% PFA, permeabilized with 0.5% Triton X-100, and stained per manufacturer protocol. Nuclei were counterstained with Hoechst 33342 (5 µg/ml). Cells were imaged by 2-channel fluorescence microscopy (Nikon Eclipse Ti) and scored using ImageJ Cell Counter [20]. Statistical significance was assessed by two-tailed Student’s t-test with variance determined by F-test.

### Immunofluorescence

Coverslip-grown astrocytes were fixed in 3.7% PFA, permeabilized with 0.5% Triton X-100, and blocked in 3% BSA/2% normal donkey serum. Primary antibodies: GFAP (Millipore AB5804, 1:500), OLIG2 (Millipore AB9610, 1:500), S100β (Abcam ab52642, 1:100), γH2AX (Novus NB100-384, 1:500), 53BP1 (Novus NB100-304, 1:500), p16^INK4a^ (Santa Cruz sc-56330, 1:200), and p53 (Santa Cruz sc-126, 1:100). Secondary antibodies: Alexa Fluor 488 and 547 (Invitrogen, donkey anti-mouse or anti-rabbit, 1:500). Nuclear stain: DAPI (2 µg/ml). Standard imaging was performed on a Nikon Eclipse Ti; confocal imaging on a Zeiss LSM 880. DNA damage foci were scored using ImageJ Cell Counter; statistics as above.

### SA-β-gal activity assay

Performed as described [21] on coverslip-grown astrocytes at 50% confluence to avoid contact inhibition. Cells were fixed, stained with X-gal substrate at pH 6.0 for 16 hours at 37°C (no CO_2_), and scored under brightfield microscopy. Statistics as above.

### DNA-damage-induced senescence

Astrocytes on coverslips were treated with 20 µM etoposide (Sigma E1383) for 3 days for DDIS development (8-week recovery in normal medium) or 100 µM etoposide for 6 hours for acute damage studies. DDIS samples for RNA-seq were treated identically (20 µM, 3 days) in proliferating P15 astrocytes, followed by 8 weeks of recovery.

### Quiescence induction

Early-passage astrocytes on coverslips were incubated for 4 days in depleted medium (0.1% AGS, 0.2% FBS, 1% P/S, 50 ng/ml BMP4; R&D Systems 314-BP) at 37°C in 3% O_2_, with fresh depleted medium every 2 days.

### DNA extraction, library preparation, and sequencing

We isolated genomic DNA using the NEB Monarch Spin gDNA Extraction Kit. Sequencing libraries were prepared following the unmodified Nanopore DNA Ligation Sequencing Kit V14 (SQK-LSK114) protocol for a Promethion flow cell. Each sample was loaded onto its own 10.4.1 PromethION Flow Cell. Sequencing was performed on a P2-solo instrument, which allows for two flow cells to be run concurrently. Two sequencing libraries were prepared for each sample. Flow cells were washed using the flow cell wash kit (EXP-WSH004-XL) midway through sequencing and a fresh sequencing library derived from the same sample was loaded. Sequencing was controlled using the MinKNOW software version 25.03.9 and was allowed to run for three days or until nearly all sequencing pores were inactive.

### RNA sequencing and analysis

RNA was extracted using TRIzol (Ambion) with on-column DNase treatment (PureLink RNA Mini Kit, Invitrogen). For the replicative senescence time course, RNA was harvested from early passage (P6, physiological O_2_) and four post-arrest time points (Sen0, Sen4, Sen8, Sen12). For oxygen comparisons, RNA was harvested from senescence-entry astrocytes in both oxygen conditions, time-matched. For DDIS, RNA was harvested from etoposide-treated cells 8 weeks after treatment. Library preparation (strand-specific, poly(A)-selected) and sequencing (Illumina HiSeq 4000, 2×150 bp) were performed by GENEWIZ. Analysis was performed on the Galaxy platform (usegalaxy.org): quality control (FastQC) [22], adapter trimming (Trimmomatic) [23], alignment to GRCh38 (HISAT2 [24], RNA-STAR [25]), gene quantification (featureCounts) [26], and differential expression (edgeR [27], DESeq2 [28]). Pathway analysis was performed using GSEA via GenePattern 2.0 [29] with KEGG and GO databases; FDR correction by Benjamini-Hochberg procedure. Heatmaps were generated using Morpheus (Broad Institute); Venn diagrams using BioVenn [30]. Custom SASP and IFN-I gene lists were as previously described [3, 14]. Ingenuity Pathway Analysis (QIAGEN) was used for additional differential expression analysis.

### RT-qPCR

RNA (500 ng) was reverse-transcribed using MultiScribe Reverse Transcriptase (Invitrogen). Quantitative PCR was performed on a ViiA 7 system using PowerUp SYBR Green master mix (Thermo Fisher). GAPDH was used as the housekeeping gene. Primer sequences are provided in Table 1. Fold-change was calculated by the 2^ΔΔCT^.

**Table 1:**
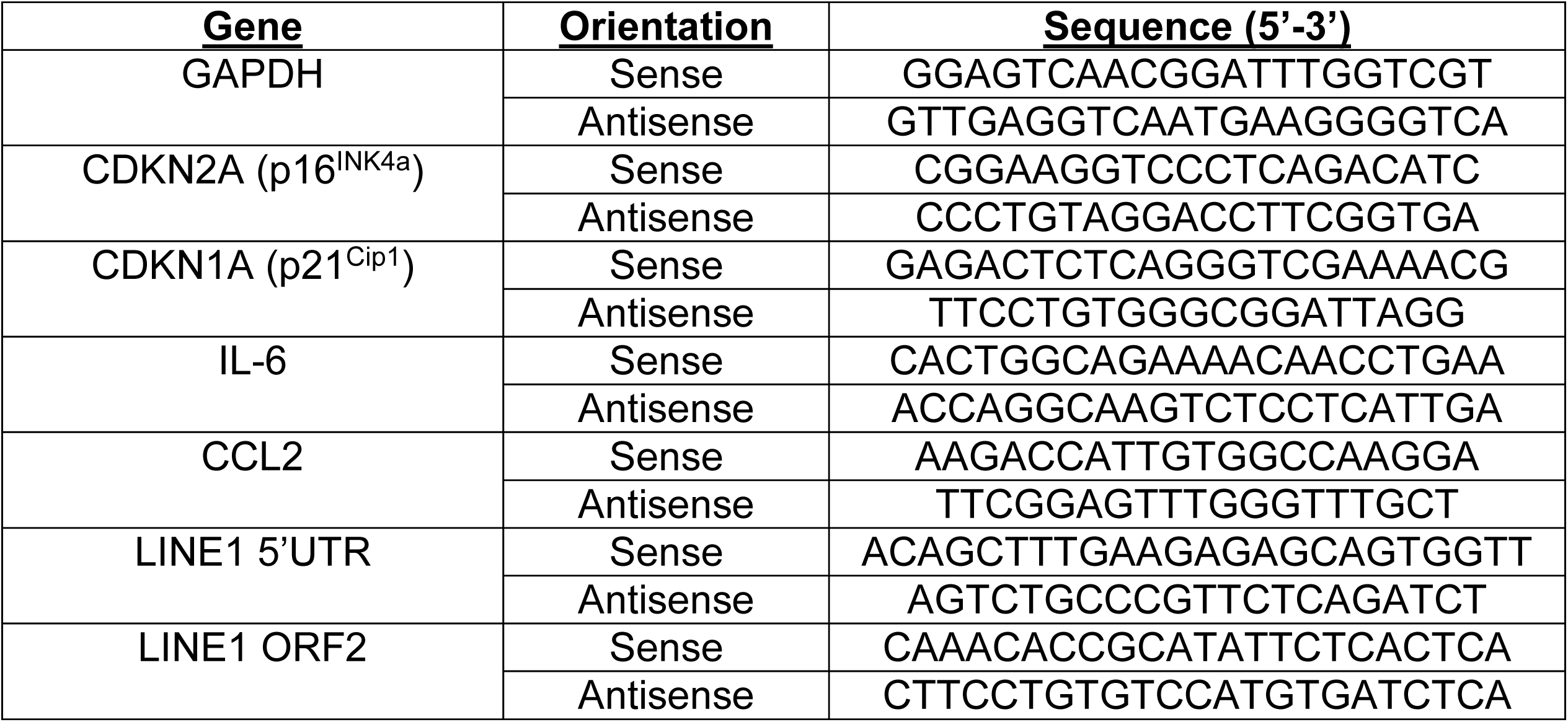
List of primer sequences for RT-qPCR. Forward and reverse primer sequences for target genes for RT-qPCR amplification. GAPDH used as housekeeping gene. Primers were designed in Ito et al., 2018, *Cell Rep*, and De Cecco et al., 2019, *Nature*.

### Nanopore basecalling and quality control

Nanopore sequencing data were basecalled using Dorado with the super-accuracy, 5mCG and 5hmCG basecalling model (dna_r10.4.1_e8.2_400bps_hac@v5.0.0_5mCG_5hmCG@v3). Basecalled reads were then aligned to the telomere-to-telomere T2T-HS1 (GCF_009914755.1) human genome [31][32]. Quality control metrics were created using PycoQC [32]. A filtered BAM to be used in various sensitive downstream applications that require only high-confidence alignments was then created. We retained only those alignments with a read quality score ≥ 10, a MAPQ ≥ 10, and filtered out secondary and supplementary alignments using samtools [33] (samtools view -b -F 2304 -q 10 -e ‘[qs] >= 10’).

### Non-reference germline insertion calling and reference genome patching

We used our TE-Seq pipeline[18] to call non-reference RTE insertions and create sample-specific, non-reference TE-patched genomes. Full methods can be found at https://doi.org/10.1186/s13100-025-00381-w. Briefly, we aligned reads to the T2T-HS1 genome using Minimap2[34] and used TLDR[35] to call non-reference insertions. Non-reference insertions were annotated as “known” or “novel” on the basis of having been identified in a database of 17 datasets of human variation; see Table S4 in Ref[35]). Putative insertions were then rigorously filtered. We required that the median MAPQ score of supporting reads be 60 (the maximum possible), that the insert have a TSD (a hallmark of retrotransposition), and that the insert have at least 10 supporting reads, three of which must fully span the insert. We then appended each insert consensus sequence, flanked by 4 kilobases (kb) of insert-site sequence to enable accurate mapping of long reads, to the T2T-HS1 genome as a standalone contig. We annotated these contigs for repetitive element content with RepeatMasker[36] and merged these annotations with the T2T-HS1 reference RepeatMasker annotations.

### LINE-1 RNA-sequencing Analysis

We used our TE-Seq pipeline[18] to analyze L1HS expression in our RNA-seq data. Full methods can be found at https://doi.org/10.1186/s13100-025-00381-w. Briefly, sequencing reads were trimmed using Fastp[37] and aligned to a sample-specific TE-patched genome (see Methods: *Non-reference germline insertion calling and reference genome patching*) using STAR[25]. Repetitive element expression was quantified using the telescope[38] tool in both “unique” and “multi” mode, which considers either uniquely mapped reads, or uniquely and multi mapped reads.

## Declarations

### Ethics approval and consent to participate

Not applicable

### Consent for publication

Not applicable

### Availability of data and materials

The TE-Seq pipeline is open-source and available at https://github.com/maxfieldk/TE-Seq. All raw and processed sequencing data have been prepared for submission and will be made publicly available in the NCBI Gene Expression Omnibus (GEO) upon peer-reviewed publication. All other supporting data are available from the corresponding author upon reasonable request.

### Authors’ contributions

Conceptualization: JMS, TAW, MMK

Data curation: TAW, MMK

Formal analysis: TAW, MMK

Funding acquisition: JMS

Investigation: TAW, MMK, JMS

Methodology: TAW, MMK, JMS

Project administration: JMS

Resources: JMS

Software: TAW, MMK

Supervision: JMS

Writing – original draft: TAW, MMK, JMS

Writing—review & editing: All authors

## Acknowledgments

We would like to thank members of the Sedivy lab and the Center on the Biology of Aging for feedback and support. We are grateful to the Brown Center for Computation and Visualization (CCV) team for their management of the OSCAR high performance computing cluster which was used throughout this work.

---

**Supplemental Figure 1:**
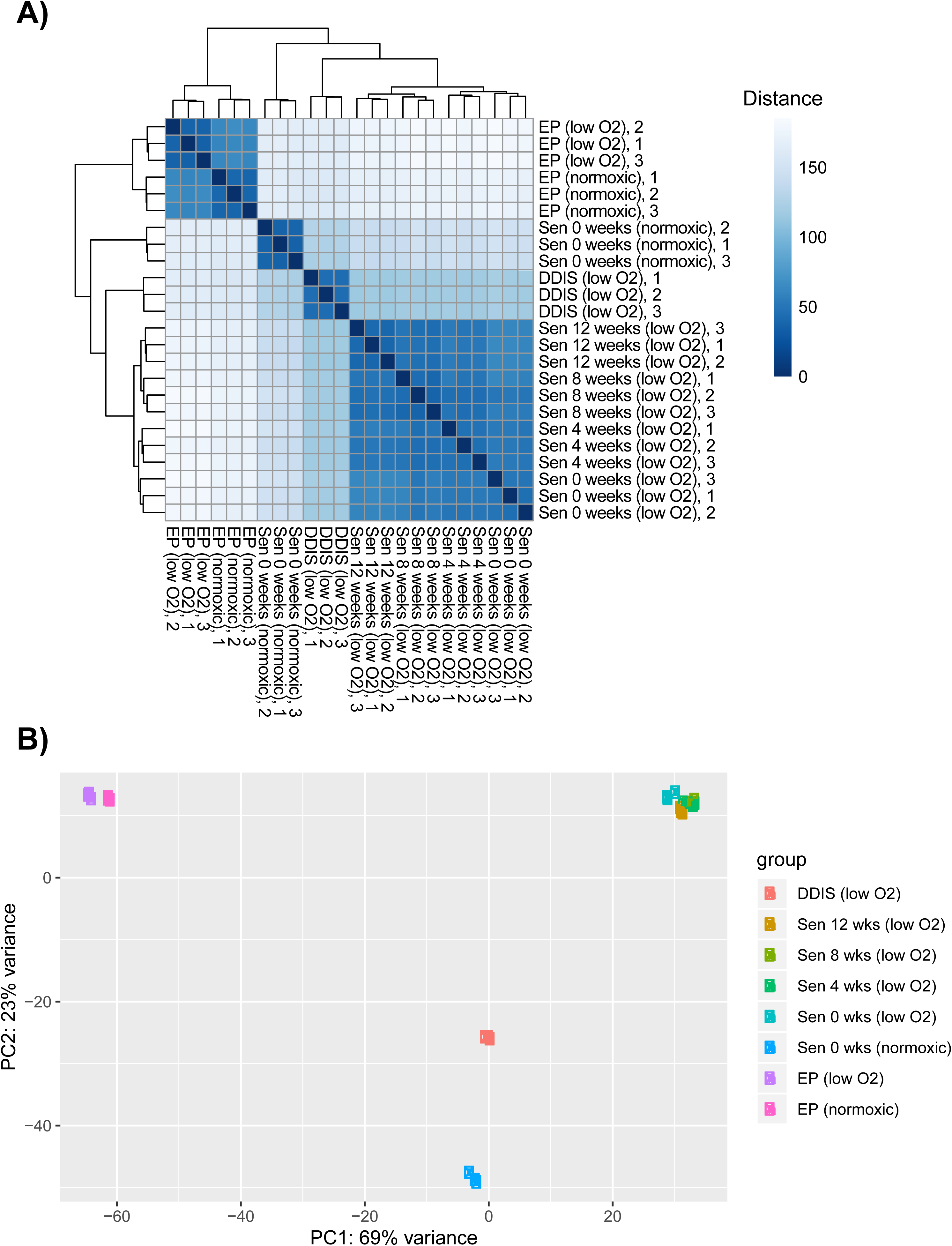
Cluster analysis for sequencing data shows conditions are different from each other while samples within a condition are similar. A) Heatmap of sample-to-sample distances with hierarchical clustering showing similarities and dissimilarities between samples. Generated by applying DESeq2 *dist* function to the transpose of the count matrix. B) PCA plot showing samples in 2D plane spanned by the first two principal components, visualizing effect of experimental covariates and batch effects. Generated using the DESeq2 *plotPCA* function.

**Supplemental Figure 2:**
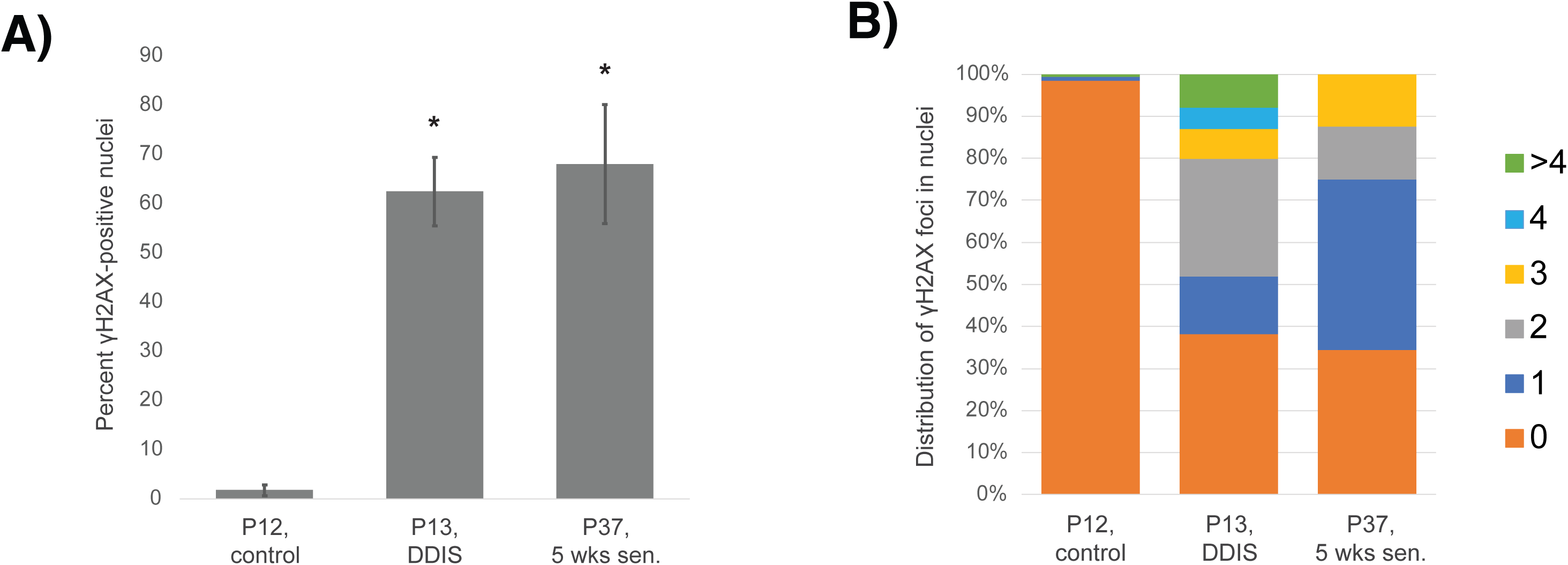
γH2AX immunofluorescent staining. Conditions imaged were early proliferative (P12), DNA-damaged (P13, DDIS), and early senescent (P37, 5 wks sen) astrocytes. Scale bars 10 µm. A) Percent positivity calculated by tabulating the number of cells with a nucleus showing at least 1 damage foci [focus - singular] as % of total nuclei. Statistics performed using a student t-test. P-value: * < 0.001. B) Distribution of damage foci in cells. Cells were scored by counting the number of foci per nucleus and sorting the cells into groups by number of foci. P12, control: 317 cells scored; P13, DDIS: 139 cells scored; P37, 5 wks sen: 32 cells scored.

**Supplemental Figure 3:**
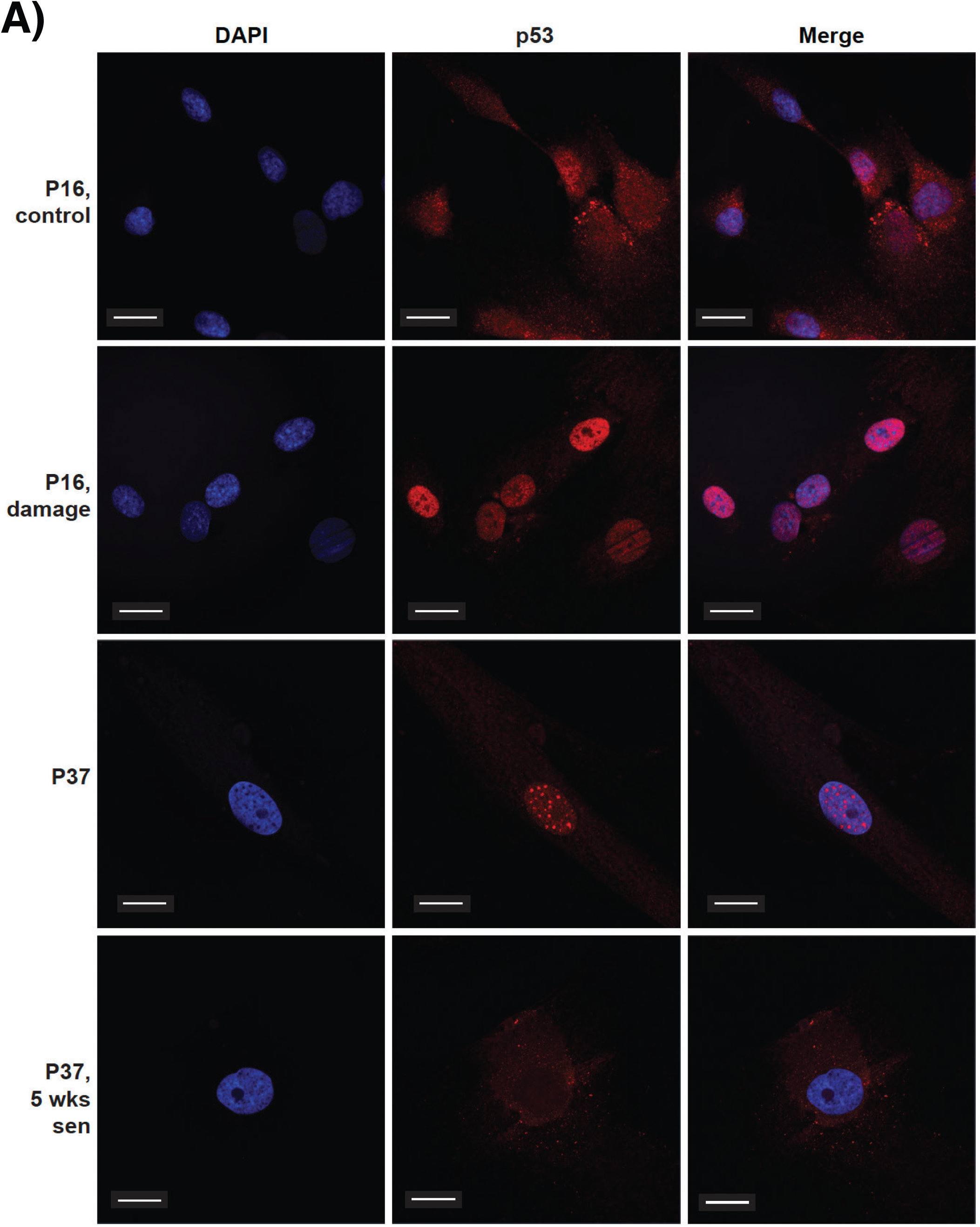
p53 immunofluorescent staining. A) Representative images of p53 immunofluorescent staining. Conditions imaged were early passage proliferative (P16, control), early passage acute damage (P16, damage), late passage arrested (P37), and late passage early senescent (P37, 5 wks sen) astrocytes. Confocal microscopy. Scale bars 20 µm.

**Supplemental Figure 4:**
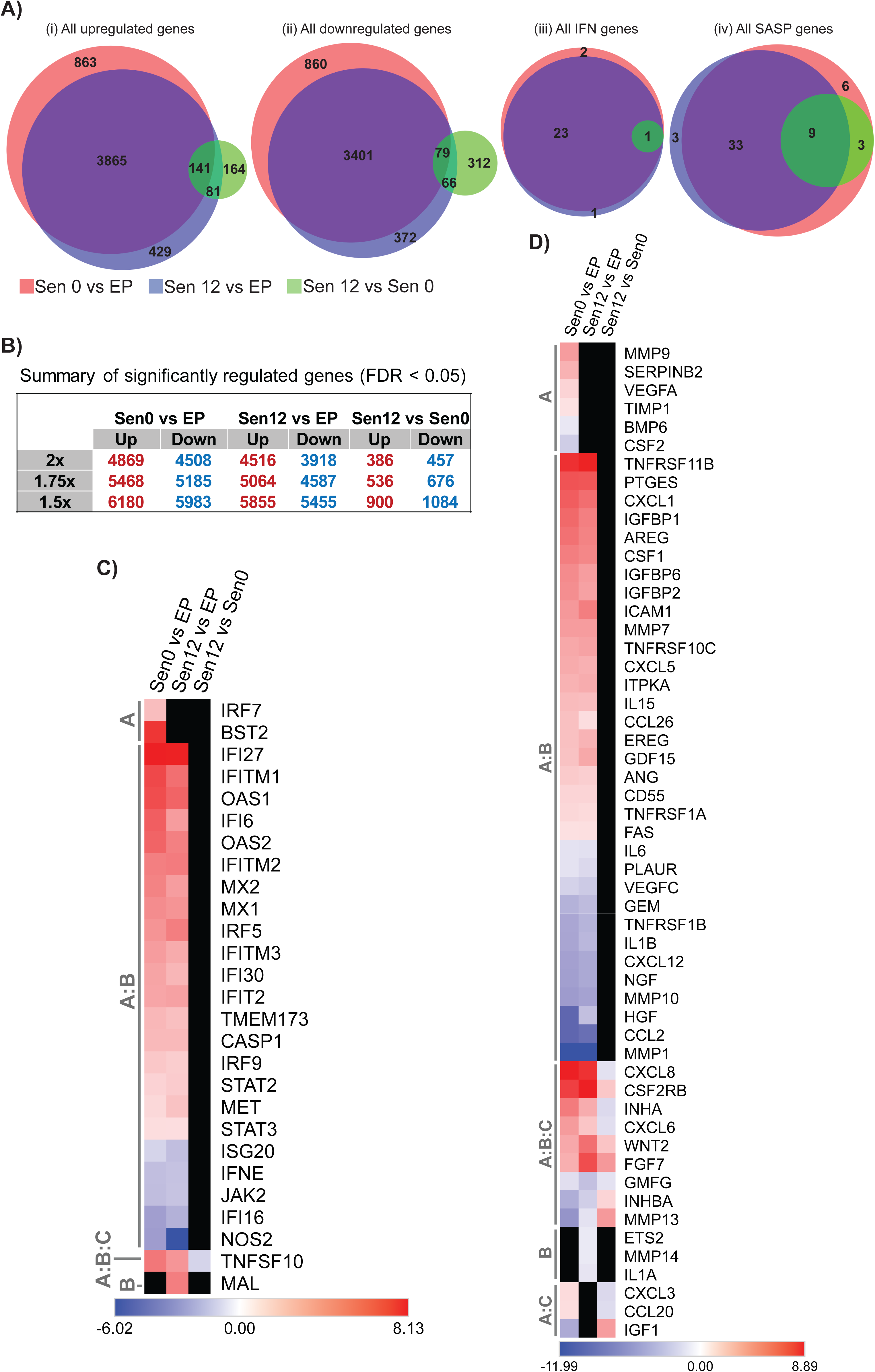
Gene-level differences in time points of replicative senescence. A) Overlap of differentially regulated genes in the comparisons of Sen0 vs. EP, Sen12 vs. EP, and Sen12 vs. Sen0 for total upregulated genes (i), total downregulated genes (ii), genes in the IFN pathway gene list (iii), and genes in the SASP pathway gene list (iv). Fold-change cutoff greater than 2, FDR cutoff less than 0.05. B) Summary of significantly regulated (FDR < 0.05) genes. Comparisons performed were Sen0 vs. EP, Sen12 vs. EP, and Sen12 vs. Sen0, for fold-change cutoffs of 1.5x, 1.75x, and 2.0x. C) Heatmap of differentially regulated IFN pathway genes for comparisons Sen0 vs. EP (Group A), Sen12 vs. EP (Group B), and Sen12 vs. Sen0 (Group C). Legend represents global log fold-change values. Fold-change cutoff 2, FDR cutoff 0.05. Black regions indicate expression values that did not meet either fold-change or statistical significance cutoffs within a comparison. D) Heatmap of differentially regulated SASP pathway genes for comparisons Sen0 vs. EP (Group A), Sen12 vs. EP (Group B), and Sen12 vs. Sen0 (Group C). Legend represents global log fold-change values. Fold-change cutoff 2, FDR cutoff 0.05. Black regions indicate expression values that did not meet either fold-change or statistical significance cutoffs within a comparison.

**Supplemental Figure 5:**
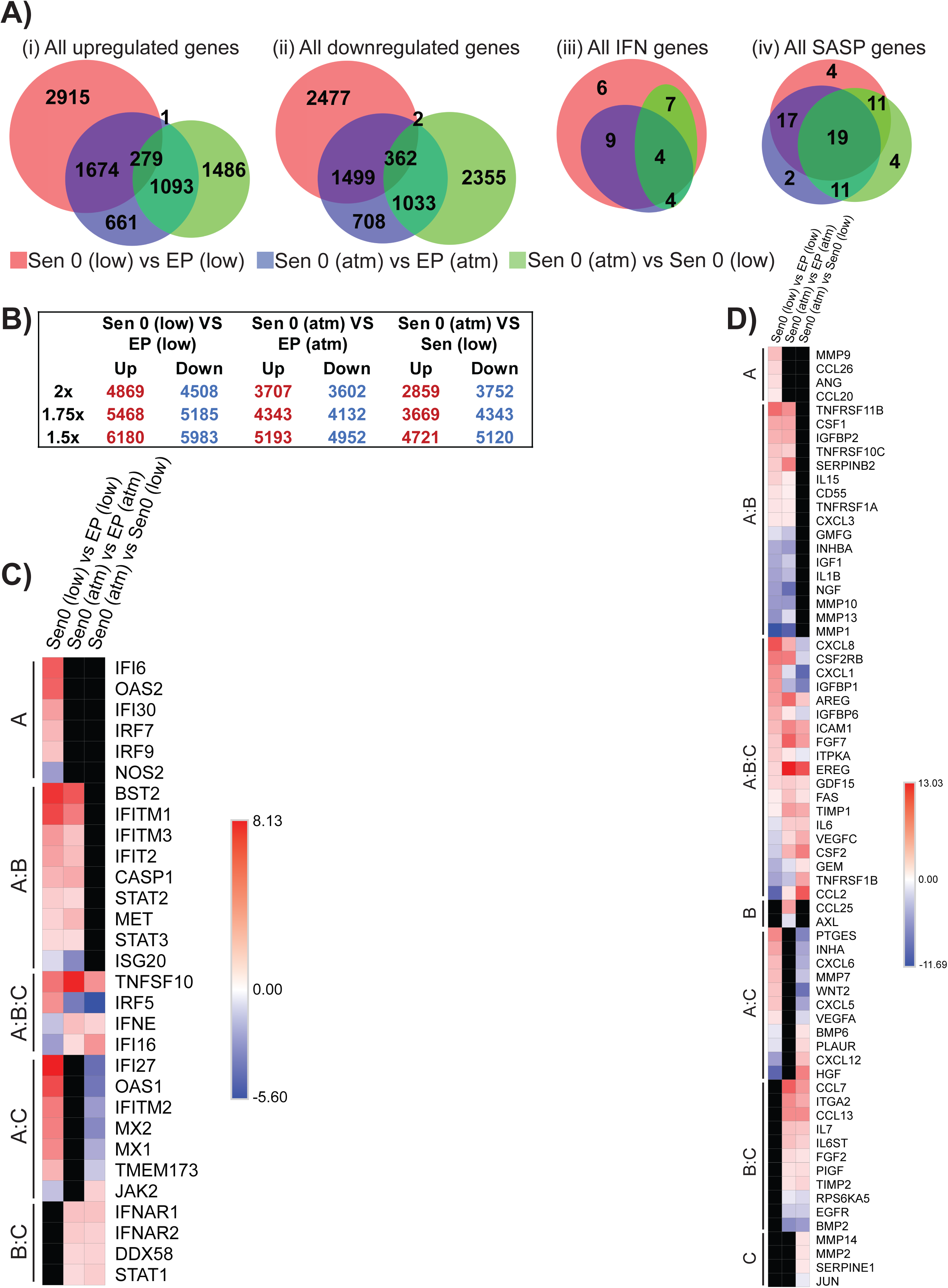
Gene-level differences in replicatively senescent astrocytes from varying oxygen culture conditions. E) Overlap of total differentially upregulated genes in the comparisons of Sen 0 (low) vs. EP (low) [red], Sen 0 (atm) vs. EP (atm) [blue], and Sen 0 (atm) vs. Sen 0 (low) [green] for total upregulated genes (i), total downregulated genes (ii), genes in the IFN pathway gene list (iii), and genes in the SASP pathway gene list (iv). Fold-change cutoff greater than 2, FDR cutoff less than 0.05. F) Summary of differentially expressed (FDR < 0.05) genes. Comparisons performed were Sen 0 (low) vs. EP (low), Sen 0 (atm) vs. EP (atm), and Sen 0 (atm) vs. Sen 0 (low), for fold-change cutoffs of 1.5x, 1.75x, and 2.0x. G) Heatmap of differentially regulated IFN pathway genes for comparisons Sen 0 (low) vs. EP (low) [A], Sen 0 (atm) vs. EP (atm) [B], and Sen 0 (atm) vs. Sen 0 (low) [C]. Legend represents global log fold-change values. Fold-change cutoff 2, FDR cutoff 0.05. Black regions indicate values that did not meet cutoffs within a comparison. H) Heatmap of differentially regulated SASP pathway genes for comparisons Sen 0 (low) vs. EP (low) [A], Sen 0 (atm) vs. EP (atm) [B], and Sen 0 (atm) vs. Sen 0 (low) [C]. Legend represents global log fold-change values. Fold-change cutoff 2, FDR cutoff 0.05. Black regions indicate values that did not meet cutoffs within a comparison.

**Supplemental Figure 6:**
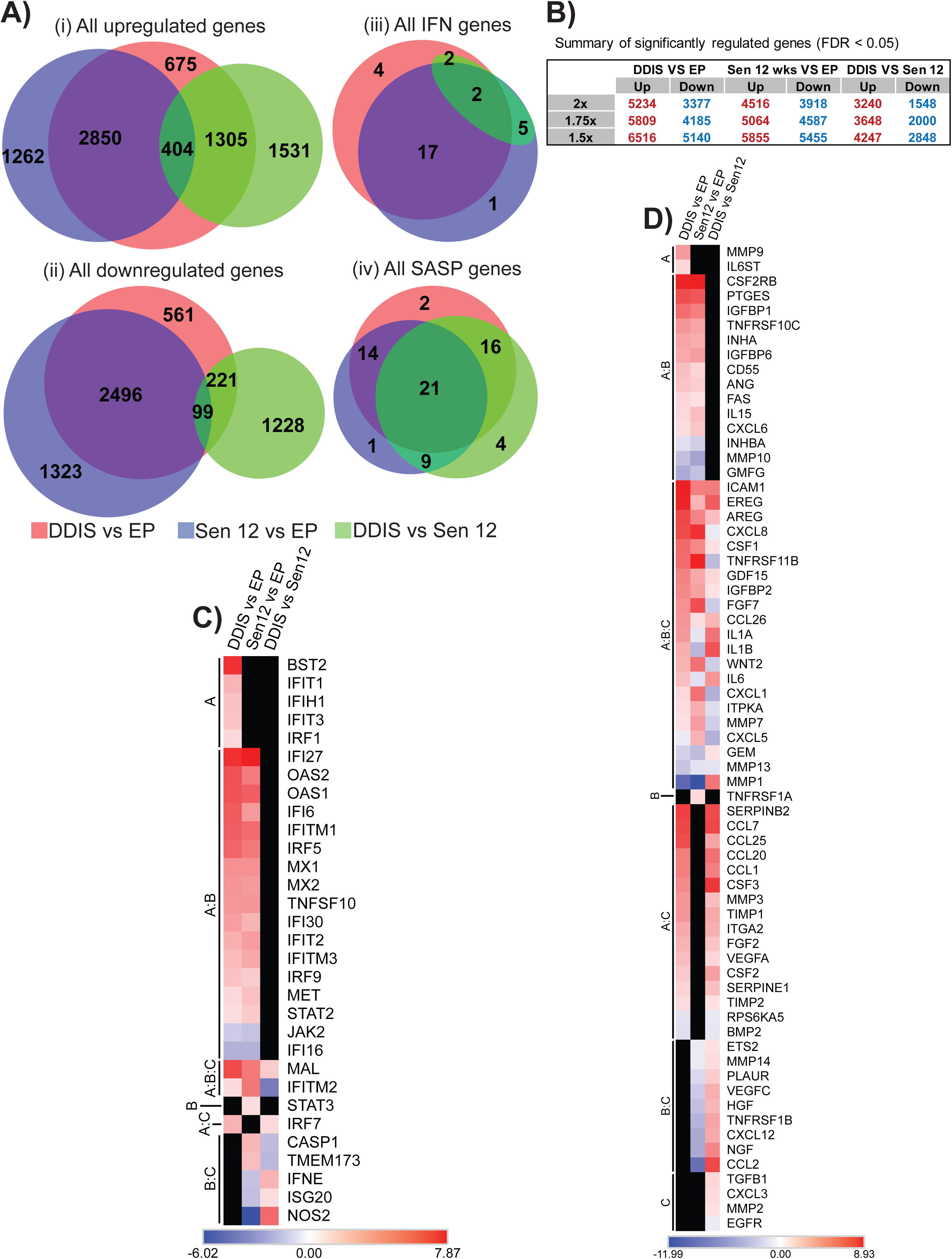
Gene-level differences in DDIS compared to replicative senescence in astrocytes. I) Overlap of total differentially upregulated genes in the comparisons of DDIS vs. EP [red], Sen12 vs. EP [blue], and DDIS vs. Sen12 [green] for total upregulated genes (i), total downregulated genes (ii), genes in the IFN pathway gene list (iii), and genes in the SASP pathway gene list (iv). Fold-change cutoff greater than 2, FDR cutoff less than 0.05. J) Summary of significantly regulated (FDR < 0.05) genes. Comparisons performed were DDIS vs. EP, Sen12 vs. EP, and DDIS vs. Sen12, for fold-change cutoffs of 1.5x, 1.75x, and 2.0x. K) Heatmap of differentially regulated IFN pathway genes for the comparisons DDIS vs. EP [A], Sen12 vs. EP [B], and DDIS vs. Sen12 [C]. Legend represents global log fold-change values. Fold-change cutoff 2, FDR cutoff 0.05. Black regions indicate expression values that did not meet cutoffs within a comparison. L) Heatmap of differentially regulated SASP pathway genes for comparisons DDIS vs. EP [A], Sen12 vs. EP [B], and DDIS vs. Sen12 [C]. Legend represents global log fold-change values. Fold-change cutoff 2, FDR cutoff 0.05. Black regions indicate expression values that did not meet cutoffs within a comparison.

**Supplemental Figure 7:**
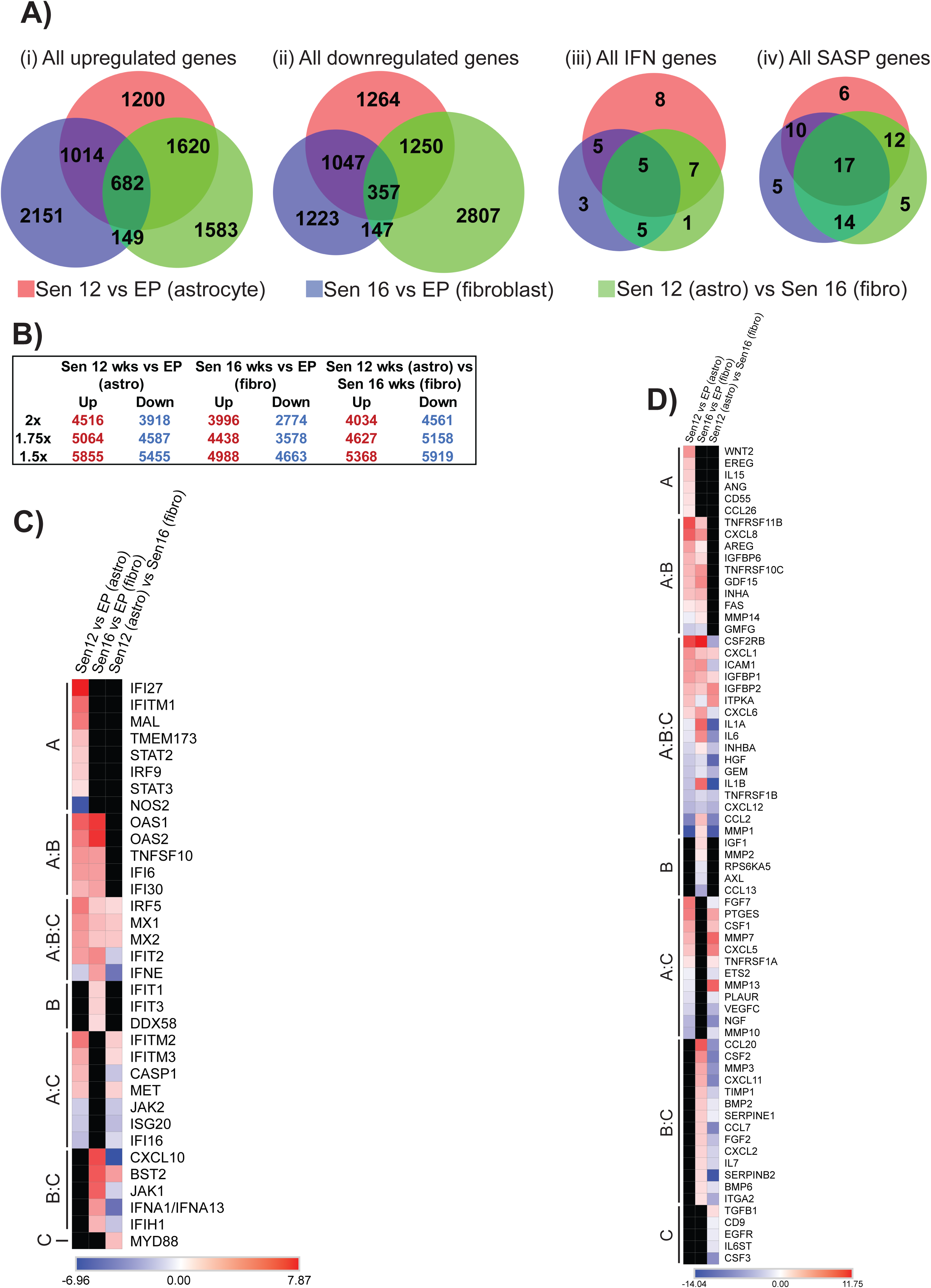
Gene-level expression changes in replicative senescence modeled in astrocytes and fibroblasts. A) Overlap of total differentially upregulated genes in the comparisons of Sen12 vs. EP (astrocytes) [red], Sen16 vs. EP (fibroblasts) [blue], and Sen12 (astro) vs. Sen16 (fibro) [green] for total upregulated genes (i), total downregulated genes (ii), genes in the IFN pathway gene list (iii), and genes in the SASP pathway gene list (iv). Fold-change cutoff greater than 2, FDR cutoff less than 0.05. B) Summary of significantly regulated (FDR < 0.05) genes. Comparisons performed were Sen12 vs. EP (astrocytes), Sen16 vs. EP (fibroblasts), Sen12 (astro) vs. Sen16 (fibro), for fold-change cutoffs of 1.5x, 1.75x, and 2.0x. C) Heatmap of differentially regulated IFN pathway genes for Sen12 vs. EP (astrocytes) [A], Sen16 vs. EP (fibroblasts) [B], and Sen12 (astro) vs. Sen16 (fibro) [C]. Legend represents global log fold-change values. Fold-change cutoff 2, FDR cutoff 0.05. Black regions indicate expression values that did not meet cutoffs within a comparison. D) Heatmap of differentially regulated SASP pathway genes for Sen12 vs. EP (astrocytes) [A], Sen16 vs. EP (fibroblasts) [B], and Sen12 (astro) vs. Sen16 (fibro) [C]. Legend represents global log fold-change values. Fold-change cutoff 2, FDR cutoff 0.05. Black regions indicate expression values that did not meet cutoffs within a comparison.

**Supplemental Figure 8:**
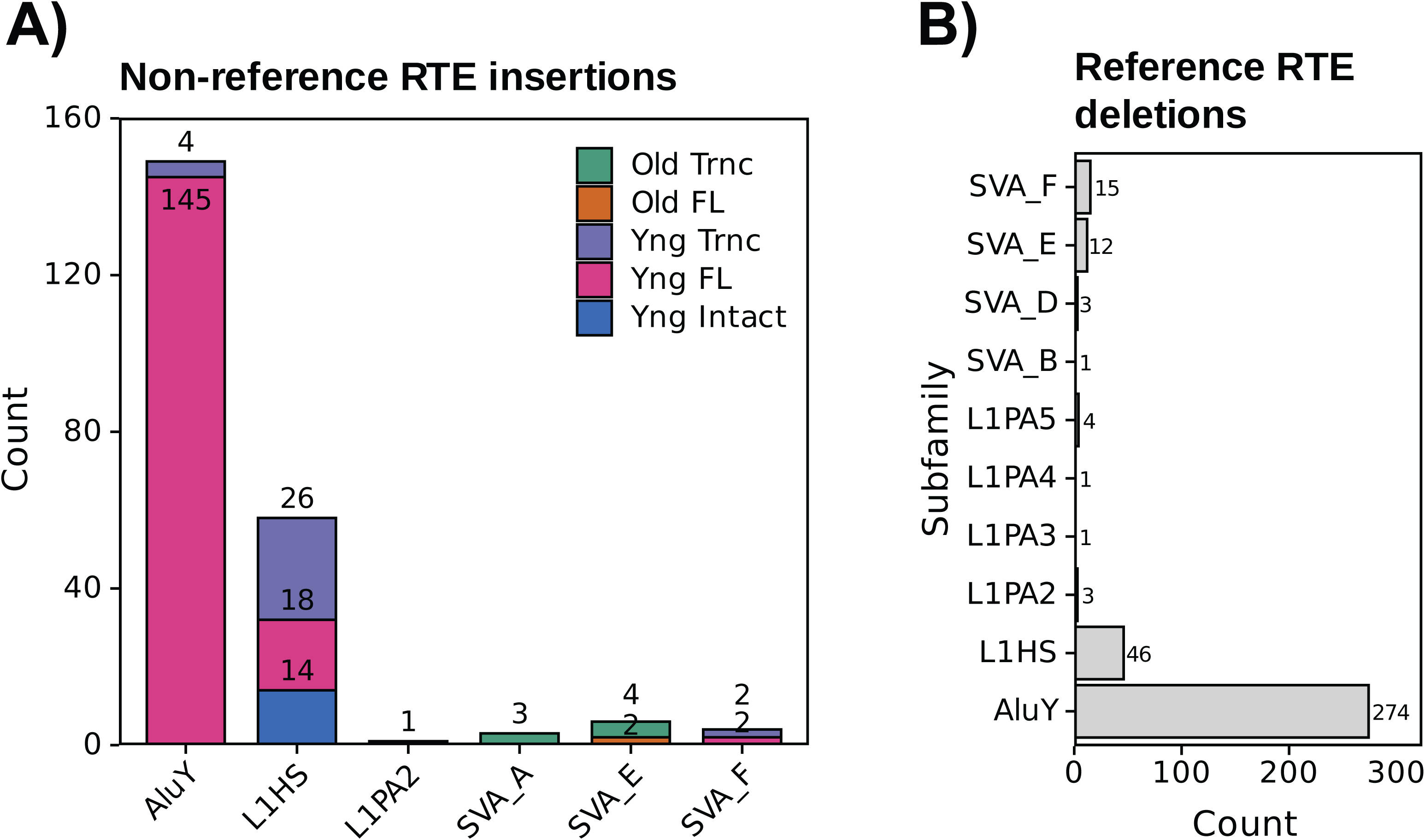
Nanopore DNA-Seq characterizes non-reference germline RTE insertions and reference RTE deletions. A) RTE insertions were called relative to the T2T-HS1 human reference genome using TLDR. Putative germline insertions were rigorously filtered (see Methods). Polymorphic germline insertion counts for L1HS, L1PA2, AluY, SVA E, and SVA F subfamilies. Insertions are colored by length, evolutionary age, and open reading frame (ORF) intactness. FL: full-length; Yng: Young; Intact: intact ORFs; Trnc: truncated. B) RTE deletions were called relative to the T2T-HS1 human reference genome using Sniffles2. Barplot shows the count of homozygous deletions, stratified by subfamily, which we masked in the custom genome.

